# Synergistic generation of cardiac resident-like macrophages and cardiomyocyte maturation in tissue engineered platforms

**DOI:** 10.1101/2024.12.04.626684

**Authors:** Meenakshi Suku, Jack F. Murphy, Sara Corbezzolo, Manus Biggs, Giancarlo Forte, Irene C. Turnbull, Kevin D. Costa, Lesley Forrester, Michael G Monaghan

## Abstract

Cardiovascular disease stands as the leading cause of death globally, claiming approximately 19million lives in 2020. On the contrary, the development of cardiovascular drugs is experiencing a decline, largely due to the bottleneck in understanding the pathophysiology of various heart diseases and assessing the effects of drugs on healthy human hearts. The development of induced pluripotent stem cell (iPSC) technology and the availability of cardiac cell types *in vitro*, has resulted in a surge in efforts to fabricate human cardiac models for disease modelling and drug discovery applications. Although numerous attempts evidence successful fabrication of 3 dimensional (3D) engineered heart tissues, the innate immune cell population of the myocardium – particularly cardiac macrophages, was until recently, overlooke. With increasing appreciation of the interactions between cardiomyocytes and macrophages in the myocardium, in this work, isogenic populations of cardiac resident-like macrophages and cardiomyocytes were generated using iPSCs, to understand the interactions between the two cell types in both 2D and 3D settings, and subjected to electric stimulation. After characterizing iPSC-derived macrophages (iMacs) and iPSC-derived cardiomyocytes (iCMs) in depth, the conditioning of iMacs to align to a cardiac resident macrophage-like phenotype in the presence of iCMs in 2D culture was explored. In co-culture with iCMs, iMacs upregulated known genes expressed by cardiac resident macrophages. Additionally, in co-culture with iMacs, iCMs displayed an elongated morphology, improved calcium function and an increase in known maturation genes such as the ratio between MYH7 and MYH6 as well as SERCA2. In a 2D setting, iMacs showed the ability to electrically couple with iCMs and facilitate synchronous beating in iCM cultures. The 2D characterisation was translated into an engineered cardiac tissue model, wherein, improvement in tissue characteristics in the presence of iMacs was demonstrated in terms of increased cell alignment, enhanced cardiomyocyte elongation, physiologically relevant beat rates and improved tissue compaction. Taken together, these findings may open new avenues to use iMacs in engineered cardiac tissue models, not only as an innate immune cell source, but also as a support cell type to improve cardiomyocyte function and maturation.

## INTRODUCTION

Cardiac diseases, ranging from arrhythmias to heart failure, represent a formidable global health burden [1], posing a significant challenge to medical science and making cardiac diseases one of the leading causes of death worldwide [2]. Until recently, our endeavours to comprehend and address these conditions were hindered by the limitations of conventional research practices, often reliant on animal models whose physiological nuances do not faithfully translate to the complexities of the human cardiovascular system. Animal cells, while valuable for certain studies, often fail to capture the intricacies of human cardiac physiology [3]. The advent of induced pluripotent stem cell (iPSC) technology has prompted significant improvement in our ability to reproduce the cellular landscape of human myocardium *in vitro* and understand their interactions with each other. *In vitro* models of the myocardium have evolved over the years, with researchers exploring various cell compositions and scaffold materials to mimic native cardiac tissue. Commonly explored *in vitro* models of the myocardium range from self-assembling organoids [4] [5], to organ-on-a-chip models [6, 7] and engineered heart and cardiac tissue models [8] [9]. While cardiomyocytes are a central component of these models, incorporating supporting cell types such as fibroblasts [10] and endothelial cells [11] has shown promise in enhancing tissue functionality.

Despite constituting a relatively small fraction of the total heart cell population, cardiac resident macrophages (CRMs) are the most abundant immune cell type in the myocardium [12], involved in governing a myriad of cellular processes including electrical conduction [13], elimination of cardiac exophers [14], facilitation of cardiomyocyte proliferation and limiting adverse remodeling post myocardial infarction (MI) [15]. In the past decade, although we have significantly improved our understanding of CRM biology and function, further refinement of our knowledge requires efficient and reliable *in vitro* culture set ups to study CRMs. Thus far, many studies have relied on blood-derived macrophages (BDMs) and immortalised cell lines [16, 17], which unfortunately do not reflect the heterogeneity of tissue resident macrophages (TRMs) [18, 19]. With limited availability of healthy human donors to study TRMs *in vitro*, novel cell types such as iPSC-derived macrophages (iMacs) have come to light as a potential candidate, as they share ontogeny with TRMs that develop in an MYB-independent, RUNX1- and SPI1-dependent pathway [20]. Importantly, previous studies have shown the potential of iMacs to mimic TRMs including microglia [21] and Kupffer cells [22] by using tissue-specific cues from conditioned media or co-culture with the primary cell type. Recent studies demonstrated the resemblance of human embryonic stem cell-derived macrophages and iMacs cultured in 3D tissues (*Biowire™ II* and engineered heart tissues respectively) to CRMs [23] so we sought to investigate the effects of distinct cardiac cues on iMacs, to derive a reliable CRM-like population *in vitro*. We exposed iMacs to conditioned media from iPSC-derived cardiomyocytes (iCMs) or co-cultured them with iPSC-derived cardiomyocytes (iCMs). After assessing the response of iMacs to these cardiac cues, we evaluated the function of iCMs in response to iMacs co-culture. We examined the impact of electrical stimulation (ES) on iCM maturation as well as iMacs polarisation. Finally, we incorporated iMacs into engineered cardiac tissues (ECTs) to verify improvement in iCM function at 3D tissue level.

## RESULTS AND DISCUSSION

### iMacs express phenotypic and genotypic differences when subjected to cardiac cues

Previous studies have modelled blood-derived macrophages (BDMs) and immortalised cell lines such as THP-1 cells to study TRMs *in vitro*. However, we chose to use iMacs as the macrophage source to model CRMs *in vitro*, due to the reported similarities of iMacs to MYB-independent yolk-sac derived TRMs [20, 22, 24]. It is critical for TRMs to maintain quiescence until triggered by an external stimulus, such as infection and pathogens [25, 26]. To assess the resting state of iMacs we compared the expression of specific genes that mark their phenotypic and functional states with BDMs. As reported in literature, iMacs were found to exhibit highly quiescent characteristics [27], with a reduced level of expression of both pro-regenerative (*TGFB*, *MRC1* and *IL10*) and pro-inflammatory genes (*NOS2*, *HIF1A* and *TNF*) compared to BDMs (Supplementary Fig.1 (a and b)). The functional capability of iMacs was confirmed by subjecting them to inflammatory triggers such as LPS and interferon gamma (IFN-γ) to obtain a pro-inflammatory response or IL-4 to yield an anti-inflammatory response as previously reported (Supplementary Fig.2). The absence of CCR2 has been used as a basic benchmark to identify CRMs in the myocardium [28]. Interestingly, iMacs expressed CCR2 at a lower level than BDMs (Supplementary Fig.1 (c)), indicating their potential as a favourable cell source to study CRMs.

Having chosen iMacs as the macrophage source, we next sought to establish the appropriate cues to educate iMacs to a more CRM-like fate *in vitro*, by either exposing iMacs to conditioned media from iCMs (iMac_CM) or by performing a co-culture of iMacs with iCMs (iMacs co-culture) (Fig. 2 (a)). Using BDMs and untreated iMacs as controls, iMacs_CM were assessed for the expression of several macrophage cell surface markers. As cardiac macrophages have been reported to express CD14 at high levels [28], the flow cytometry analysis performed in this study followed a similar gating strategy to previous work by Bajpai *et al.* [28], thus gating for cell populations expressing CD14. In comparison to BDMs, iMacs and iMacs_CM expressed almost 41% more CD14, highlighting a phenotype more akin to TRMs [29]. Compared to untreated iMacs, iCM conditioned media treatment was found to increase the CD14^+^CCR2^+^ population by 67% (Fig. 2 (d)). Upregulation of CCR2 expression was confirmed using immunofluorescence imaging. In line with earlier observations, iMacs were negative for CCR2, while BDMs and iMacs_CM were found to express comparable levels (Fig. 2 (b and c)). CCR2 upregulation is typically linked to chemotactic and migratory responses in macrophages [30]. Since CRMs are known to be CCR2^-^ with high CD14 expression [28], further evaluation of iMacs_CM was performed on the CD14^+^CCR2^-^ cell populations. The expression of CX3CR1, which has been previously shown to be abundant in murine CRMs [31] [32], was found to be significantly upregulated in CD14^+^CCR2^-^ iMacs_CM by almost 60% in comparison to untreated iMacs (Fig. 2 (e)). Additionally, the expression of HLA-DR, the human equivalent to MHCII, was 25% downregulated in iMacs_CM in comparison to iMacs (Fig. 2 (f)). Regardless, iMacs_CM were functionally active, showing comparable phagocytic and polarisation potential to iMacs (Supplementary fig. 3). Notably, the response of iMacs_CM to LPS treatment was found to highly suppressed in comparison to LPS treated iMacs, with a 97% downregulation in IL6 release, 91% downregulation in TNFα release and 95% downregulation in IL10 release (Supplementary fig. 3). Further functional assessment was performed on iMacs_CM using immunofluorescence imaging. Both iMacs and iMacs_CM expressed connexin-43 (Cx43), with iMacs_CM displaying a 35% upregulation in the expression (Fig. 2 (g and h)).

**Fig. 1.**
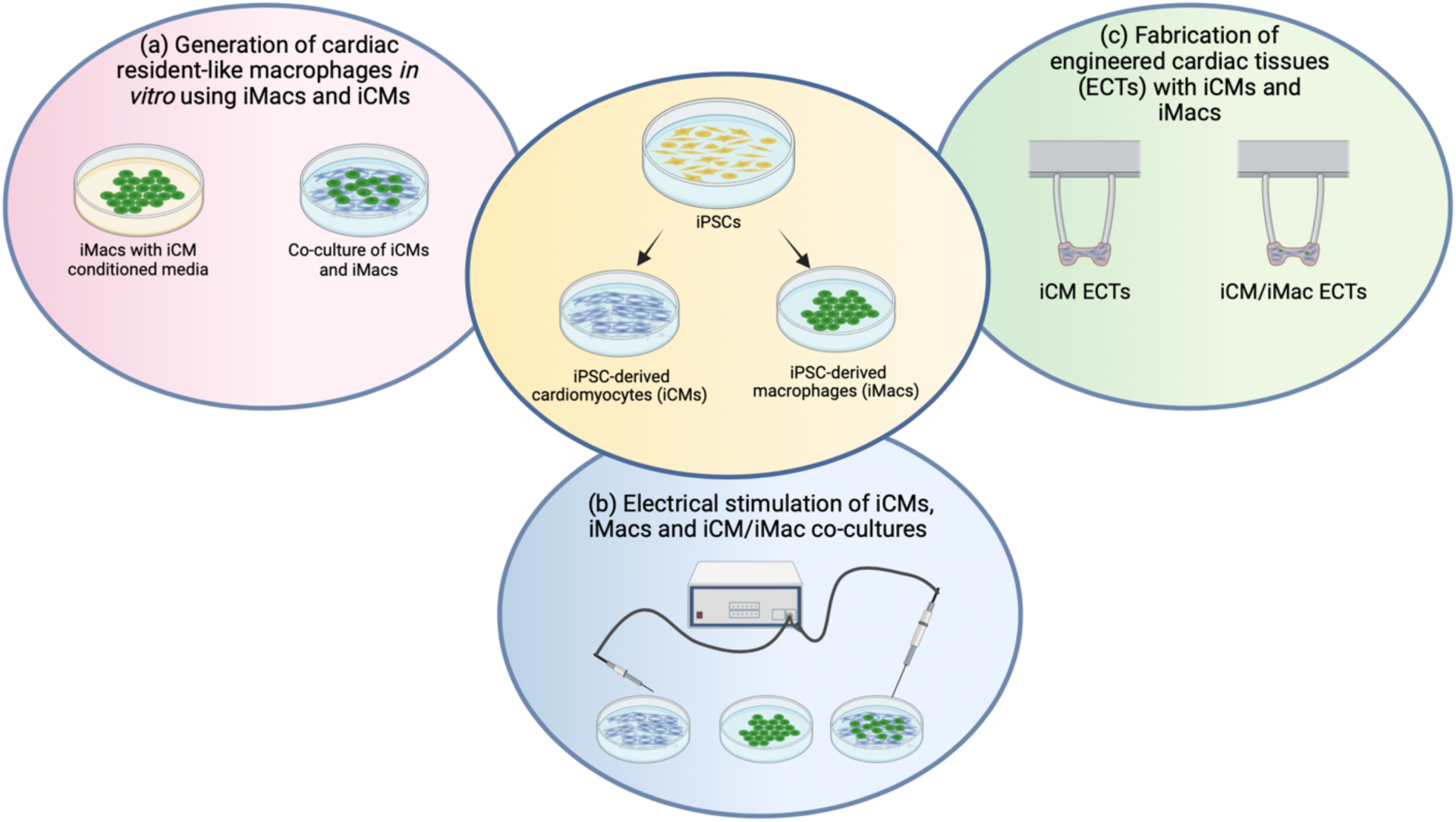
Graphical abstract. This study first delves into two potential strategies to generate cardiac resident-like macrophages *in vitro*, by exposing iPSC-derived macrophages (iMacs) to isogenic iPSC-derived cardiomyocyte (iCM) conditioned media, or by performing co-cultures of iMacs and iCMs. Next, the effect of electrical stimulation on iCMs, iMacs and iCM/iMac co-cultures is explored. Finally, the benefits of iMac addition into iCM cultures is demonstrated in a 3D engineered cardiac tissue model.

**Fig. 2.**
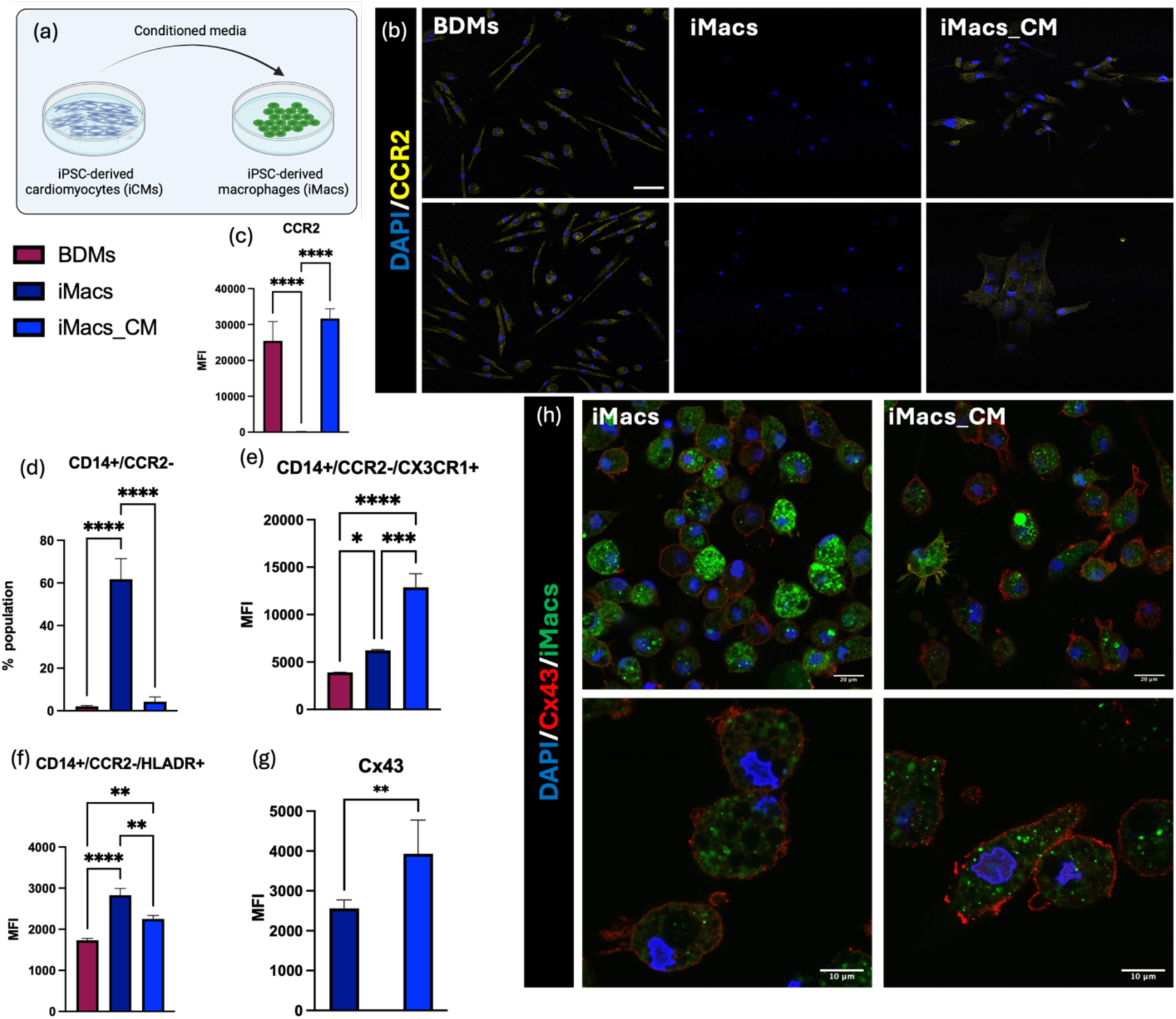
iMacs exhibit phenotypic and functional differences when subjected to iCM conditioned media. (**a**) Schematic representation of the experimental setup, (**b**) Representative images of BDMs, iMacs and iMacs_CM stained with DAPI and CCR2. Scale bar – 100 µm, (**c**) Mean fluorescence intensity (normalised to cell number) of CCR2 expression, quantified using ImageJ, (**d**) Quantification of CD14^+^CCR2^-^ populations of BDMs, iMacs and iMacs_CM. (**e**) Quantification of CD14^+^CCR2^-^CX3CR1^+^ populations of BDMs, iMacs and iMacs_CM. (**f**) Quantification of CD14^+^CCR2^-^HLA-DR^+^ populations of BDMs, iMacs and iMacs_CM. (**g**) Mean fluorescence intensity (normalised to cell number) of connexin43 expression, quantified using ImageJ, (**h**) Representative images of iMacs and iMacs_CM stained with DAPI and Connexin43. Scale bar – 20 µm, magnified image scale bar – 10 µm. Statistical analysis - one-way ANOVA with Tukey’s comparison test for c, d, e, and f; Student T test with Welch’s correction for g. * represents P<0.05, ** represents P<0.01, *** represents P<0.001 and **** represents P<0.0001. MFI – mean fluorescence intensity.

### iMacs display a more CRM-like phenotype when in co-culture with iCMs

With encouraging responses in iMacs following treatment with iCM-conditioned media, the subsequent goal was to investigate how iMacs interact with iCMs in a co-culture setting. In murine hearts, the ratio of cardiomyocytes to CRMs have been reported to be 1:5 [14]. However, in this study, a 5:1 ratio of iCMs to iMacs was used when performing co-cultures (Fig. 3 (a)) based on an average reported by Litviňuková *et al.* using single-cell analysis of adult human atrial and ventricular heart tissue [33]. Despite iCMs being plated at high cell densities (200,000 cells/cm^2^), iMacs were found to integrate well into iCM cultures (Fig. 3 (b)), notably into beating patches of iCMs (Supplementary video 1). As previously demonstrated, both iCMs and iMacs express Cx43. In the myocardium, cardiomyocytes and CRMs communicate with each other through Cx43 gap junctions [13], and hence, the expression of Cx43 was assessed. Although shared Cx43 gap junctions could not be isolated in iCM/iMacs co-cultures, both the cell types abundantly expressed Cx43 (Supplementary fig. 4). Additionally, to examine potential pro-inflammatory activation of iMacs caused due to iCM media (RPMI media with B27™), secretome from iMacs_CM and iMacs_co-culture were collected and ELISAs were performed for the presence of pro-inflammatory cytokines. Untreated iMacs and iMacs cultured in RPMI media with B27™ (supplemented with 100 ng/ml M-CSF; iMacs_RPMI+B27) were used as controls. TNFα and IL10 productions were found to be similar in all culture conditions (Supplementary fig. 5). However, over 60% more IL6 was found to be present in iMacs_CM and iMacs_RPMI+B27 secretome, potentially indicating mild activation (Supplementary fig. 5). Notably, iMacs displayed quiescence when in co-culture with iCMs, with strong suppression in their IL6 release (Supplementary fig. 5).

**Fig. 3.**
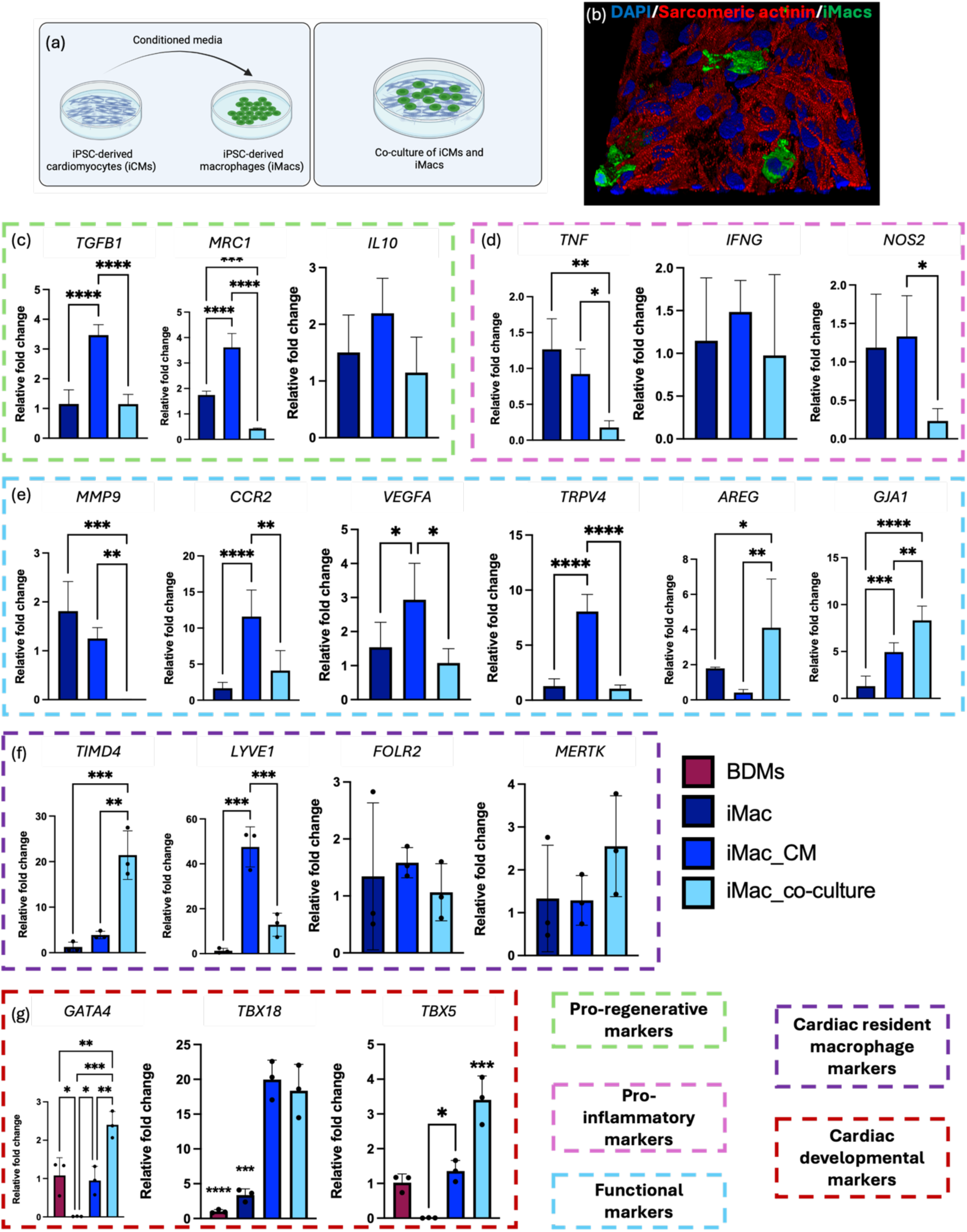
iMacs display a more CRM-like phenotype when in co-culture with iCMs. (**a**) Schematic representation of the experimental setup, (**b**) Representative 3D reconstructed image of iCMs stained with Sarcomeric actinin, highlighting the integration of iMacs into iCM cultures. iCMs generated using SFCi55 iPSC line and iMacs generated using zombie green tagged SFCi55 iPSC line. Scale bar – 50 µm, (**c**) Expression of a set of pro-regenerative genes quantified using qPCR, (**d**) Expression of a set of pro-inflammatory genes, (**e**) Expression of a set of functional genes, (**F**) Expression of a set of cardiac resident macrophage-specific genes, (**f**) Expression of a set of cardiac developmental genes. Statistical analysis - one-way ANOVA with Tukey’s comparison test. * represents P<0.05, ** represents P<0.01, *** represents P<0.001 and **** represents P<0.0001. Fold change of mRNA relative to iMacs for (c, d, e and f) and to BDMs for (g).

In attempt to further characterise iMacs_CM and iMacs_co-culture, their expression of a panel of genes was assessed, using untreated iMacs as control. While iMacs_CM displayed a highly upregulated pro-regenerative response, with a three-fold increase in *TGFB1* and a two-fold increase in *MRC1* expressions, iMacs_co-culture maintained quiescence, with no difference in their expression of *TGFB1*, *MRC1* and *IL10* in comparison to iMacs (Fig. 3 (c)). On the other hand, the expression of pro-inflammatory genes such as *TNF*, *IFNG* and *NOS2* were comparable between iMacs and iMacs_CM, while iMacs_co-culture exhibited a 90% lower level of TNF and 80% lower level of *NOS2* compared to iMacs (Fig. 3 (d)). Notably, previous work by Li *et al.* to study the interaction between cardiomyocytes and macrophages *in vitro* (using HL-1 and RAW264.7 cell lines), showed a reduced level of expression of pro-inflammatory markers when macrophages are exposed to cardiomyocyte conditioned media [34]. Taken together, these results highlight the differences in the activation states of iMacs_CM and iMacs_co-culture, with iMacs_co-culture displaying a quiescent and anti-inflammatory phenotype (Fig. 3 (c and d)). In addition to the activation states, the functional ability of macrophages is also of great importance. Matrix metalloproteinase-9 (*MMP9*) plays a direct role in ECM degradation and activation of cytokines and chemokines, for the regulation of tissue remodeling [35]. Furthermore, in a number of pathological conditions such as arthritis, diabetes and cancer, high *MMP9* production is highlighted as a biomarker for inflammation [36]. In the context of cardiovascular diseases, the deletion of *MMP9* has been proven to be beneficial, to facilitate effective tissue remodeling post-MI [37]. In this study, the expression of *MMP9* was found to be 99% lower in iMacs_co-culture compared to iMacs (Fig. 3 (e)), indicating homeostasis and a potential for robust matrix production [35]. The expression levels of *CCR2*, *VEGFA* and *TRPV4* further highlighted the quiescence of iMacs_co-culture (Fig. 3 (e)). In iMacs_CM the expression of *CCR2* transcript was consistent with the protein expression (Fig. 2 (b, c and d)), with an 85% increase compared to iMacs (Fig. 3 (e)). Remarkably, iMacs_co-culture maintained the CCR2^low^ phenotype, which is of high significance, much like CRMs [13, 14, 27, 28, 31, 38–43]. In the myocardium, CCR2^+^ and CCR2^-^ macrophages express TRPV4 in abundance [44]. While *TRPV4* and the angiogenesis marker *VEGFA* expressed at significantly higher levels (84% and 48% respectively) in iMacs_CM, the expression level of these genes was comparable between iMacs_co-culture and iMacs (Fig. 3 (e)). Although the quiescence of iMacs_co-culture was evident in most of the genes assessed, their functional potential was highlighted with the expressions of *AREG* and *GJA1* (Fig. 3 (e)). As previously mentioned, CRMs couple with cardiomyocytes *in vivo* through Cx43 gap junctions [13]. Dephosphorylation of Cx43 has been shown to affect cardiac function, by damaging the electrical coupling [45]. Additionally, more recent studies have demonstrated the role of amphiregulin (AREG), a growth factor produced by CRMs, in carrying out efficient Cx43 phosphorylation [46]. A 58% upregulation in *AREG* and 84% upregulation in *GJA1* (gene encoding Cx43) was found in iMacs_co-culture when compared to iMacs (Fig. 3 (e)). Collectively, these findings highlight the quiescent state of iMacs_co-culture, with a steady downregulation in *CCR2* expression and upregulation in *AREG* and *GJA1* expression, demonstrates an appropriate interaction between iMacs and iCMs.

According to most recent studies, macrophage populations in the myocardium include TLF^+^ macrophages which are maintained through self-renewal with minimal monocyte input (expressing TIMD4 and/or LYVE1 and/or FOLR2), CCR2^+^ macrophages that are almost entirely replaced by monocytes (TIMD4^−^LYVE1^−^FOLR2^−^), and MHC-II^hi^ macrophages that are partially replaced by circulating monocytes (TIMD4^−^LYVE1^−^FOLR2^−^CCR2^−^) [42]. Observing pronounced differences in the expression of CCR2 between iMacs_CM and iMacs_co-culture (iMacs_CM being CR2^high^ and iMacs_co-culture being CCR2^low^), the expression of other known CRM markers such as TIMD4, LYVE1 and FOLR2 was analysed. While iMacs_co-culture showed a 94% increase in TIMD4 expression, iMacs_CM demonstrated a 97.2% increase in LYVE1 expression, compared to iMacs (Fig. 3 (f)). In comparison to iMacs, iMacs_co-culture also exhibited an 89% increase in LYVE1 expression (Fig. 3 (f)). FOLR2 and MERTK were expressed at a comparable level in all three groups (Fig. 3 (f)). The expression of MERTK was also found to be comparable across the three groups, with a subtle trend of upregulation in iMacs_co-culture (Fig. 3 (f)).

During the process of differentiating iMacs, non-adherent monocyte-like macrophage precursors (pre-macs) are collected and matured for 7 days to generate functional iMacs. Pre-macs generated using this protocol are considered ideal precursors for the generation of macrophages resembling key features of TRMs such as alveolar macrophages or microglia cells [20, 47–49]. Bearing this in mind, iMacs_CM and iMacs_co-culture were assessed for the expression of cardiac developmental markers, to understand the impact of cardiac cues on iMacs. BDMs were used as the control, to make comparisons. It is important to note that, iMacs_co-culture expressed a 99.2% increase of GATA4, 71% increase of *TBX18* and 99.9% increase of *TBX5*, in comparison to iMacs as assessed by qPCR (Fig. 3 (g)). Additionally, the *GATA4* and *TBX5* expression in iMacs_co-culture was 43.4% and 51% (respectively) more than iMacs_CM (Fig. 3 (g)). Taken together, the expression of all genes analysed show that iMacs respond differently to being exposed to iCM conditioned media and to being in physical contact with iCMs. In particular, the responses of iMacs in co-culture with iCMs demonstrate a profile that resembles CRMs, with a downregulation in the expressions of *MMP9 and CCR2*; and an upregulation in the expressions of *AREG, GJA1, TIMD4, LYVE1, GATA4, TBX18 and TBX5*.

### iCMs display signs of improved maturation when in co-culture with iMacs

Following analyses of the phenotypic and genotypic characteristics of iMacs in co-culture with iCMs, it was of interest to comprehend the response of iCMs to the presence of iMacs in co-culture (Fig. 4 (a)). In adult hearts, cardiomyocytes exhibit an elongated and rod-shaped morphology, with approximately 65% of them being mononucleated—a percentage that remains relatively constant throughout life [50, 51]. Despite the current advancements in differentiation efficiency, iCMs continue to display a smaller size and a rounder shape, indicating an immature or foetal-like phenotype [52]. By performing immunofluorescence imaging for cardiac troponin T (cTnT) on iCM/iMacs co-cultures, a clear difference was noted in the phenotype of iCMs in co-culture with iMacs, with an elongated morphology, comparable to adult cardiomyocytes (Fig. 4 (b)). On quantifying the elongation using aspect ratio which is the ratio between the long axis to the short axis of a cell with perfect circle has an aspect ratio of 1 and elongated cells have an aspect ratio greater than 1. iCMs in co-culture with iMacs displayed a clear and significant 58.7% increase in the aspect ratio (Fig. 4 (c)) and this elongated morphology was also observed using Sarcomeric actinin staining (Fig. 4 (d)).

**Fig. 4.**
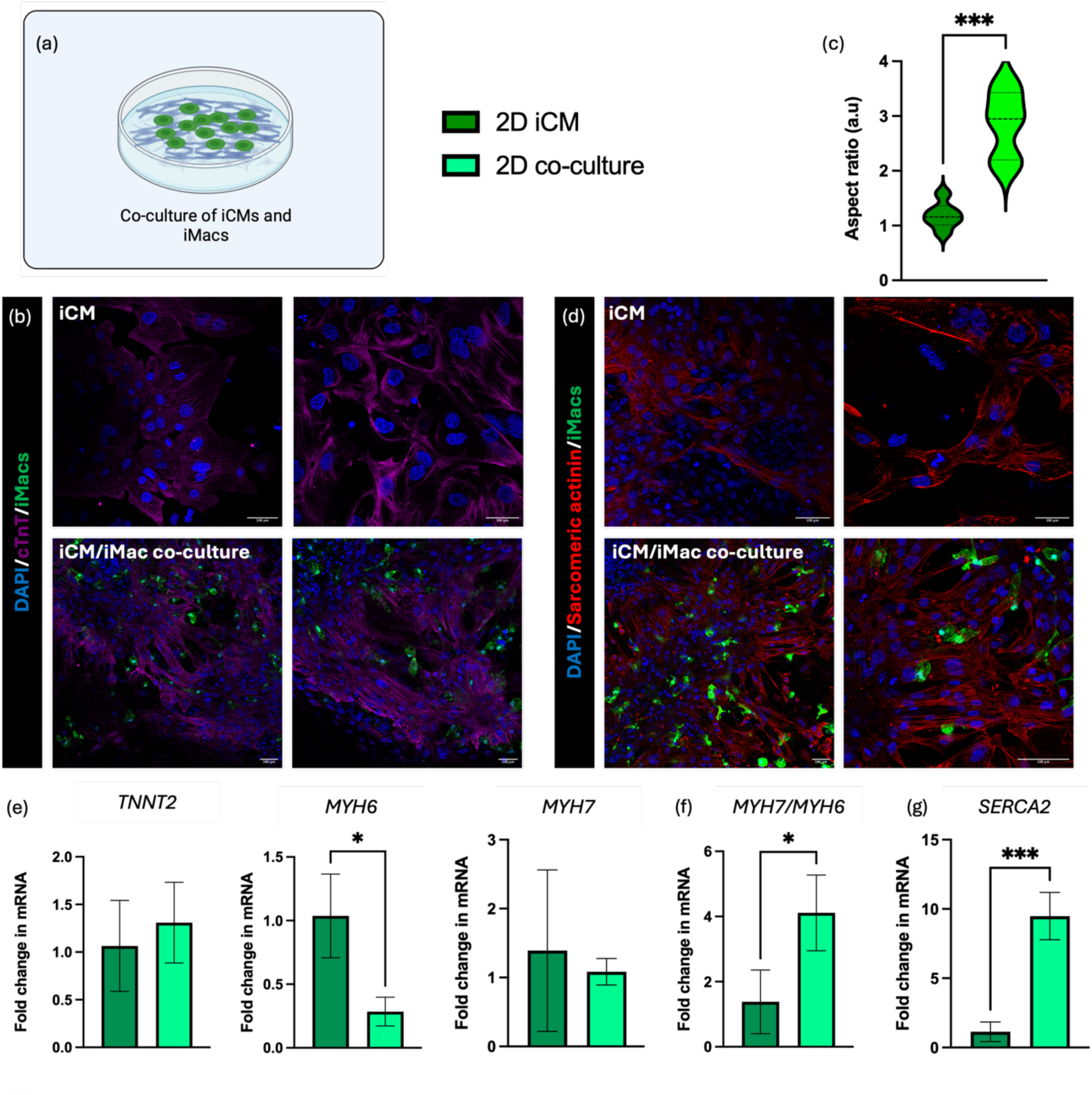
iCMs are more elongated, with improved expression of maturation genes, when in co-culture with isogenic iMacs. (**a**) Schematic representation of the experimental setup, (**b**) Representative images of iCMs and iCM/iMacs co-cultures stained with cardiac troponin T (cTnT). iCMs generated using SFCi55 iPSC line and iMacs generated using zombie green tagged SFCi55 iPSC line. Scale bar – 100 µm (**c**) Quantification of cell elongation by measuring aspect ratio of iCMs, (**d**) Representative images of iCMs and iCM/iMacs co-cultures stained with Sarcomeric actinin. Scale bar – 100 µm, (**e**) Expression of a set of cardiomyocyte-specific genes, using qPCR. (**f**) Ratio of the expression of *MYH7*/*MHY6*, using qPCR. (**g**) Expression of *SERCA2*, using qPCR. Statistical analysis - Student T test with Welch’s correction for c, e f, and g. * represents P<0.05 and *** represents P<0.001.

To investigate improvement in iCM maturity, the expression of several genes was assessed. The gene encoding cTnT, *TNNT2*, was found to be expressed at comparable levels in iCMs and iCMs in co-culture with iMacs (2D co-culture) (Fig. 4 (e)), indicating no dedifferentiation of iCMs. Other common markers for cardiomyocyte maturation included *MYH6* (encoding α-MHC; predominant MHC in the adult atrium [53, 54]) and *MYH7* (encoding β-MHC; predominant MHC in the adult ventricles [53]). In humans, the ratio of *MYH7*/*MYH6* has been shown to increase with age [55], as the levels of *MYH6* decreases [56]. This trend is also reported in extended culture conditions of iCMs [57] as well as in engineered heart tissues [58]. In comparison to iCMs, iCMs in co-culture with iMacs expressed comparable levels of MYH7 and 70% less *MYH6* (Fig. 4 (e)). Significantly, the *MYH7*/*MYH6* ratio was found to be 62.5% higher in iCMs that were in co-culture with iMacs, demonstrating a more mature phenotype (Fig. 4 (f)). Another key function of cardiomyocytes, is their ability to handle calcium efficiently, and sarcoplasmic/endoplasmic reticulum calcium ATPase (SERCA) is a key buffer responsible for calcium uptake preceding diastole [59]. Remarkably, a nine-fold (88.8%) increase in expression of *SERCA2* was noted in iCMs in co-culture with iMacs, further highlighting their maturation in terms of gene expression (Fig. 4 (g)). Collectively, these findings indicate improvement in iCM phenotype and maturation, when in co-culture with iMacs.

### iMacs electrically couple with iCMs and facilitate synchronous beating

To further characterise the functional aspects of iCM maturation, their intracellular calcium handling properties was evaluated using Cal520^®^ AM dye (Fig. 5 (a)). Asynchronous and heterogeneous spontaneous beating rates is one of the well-documented shortcomings of iCMs [60, 61]. In an attempt to synchronise the beating of iCMs, previous studies have used a number of different approaches, such as electrical stimulation [62] [63] or fabrication of 3D tissue constructs [64], using mechanical stimulation approaches [60, 65]. On assessing nine different regions of interest within a well of iCMs, there was no notable synchrony in the beating of iCMs (Fig. 5 (b)). However, on performing a similar assessment of iCM/iMacs co-cultures, all the nine regions appeared to beat in synchrony, with peaks corresponding to iCM contractions (Fig. 5 (b)); synchrony highlighted using red lines). This suggests that the addition of iMacs within iCM cultures could be an excellent solution to facilitating synchronous beating.

**Fig. 5.**
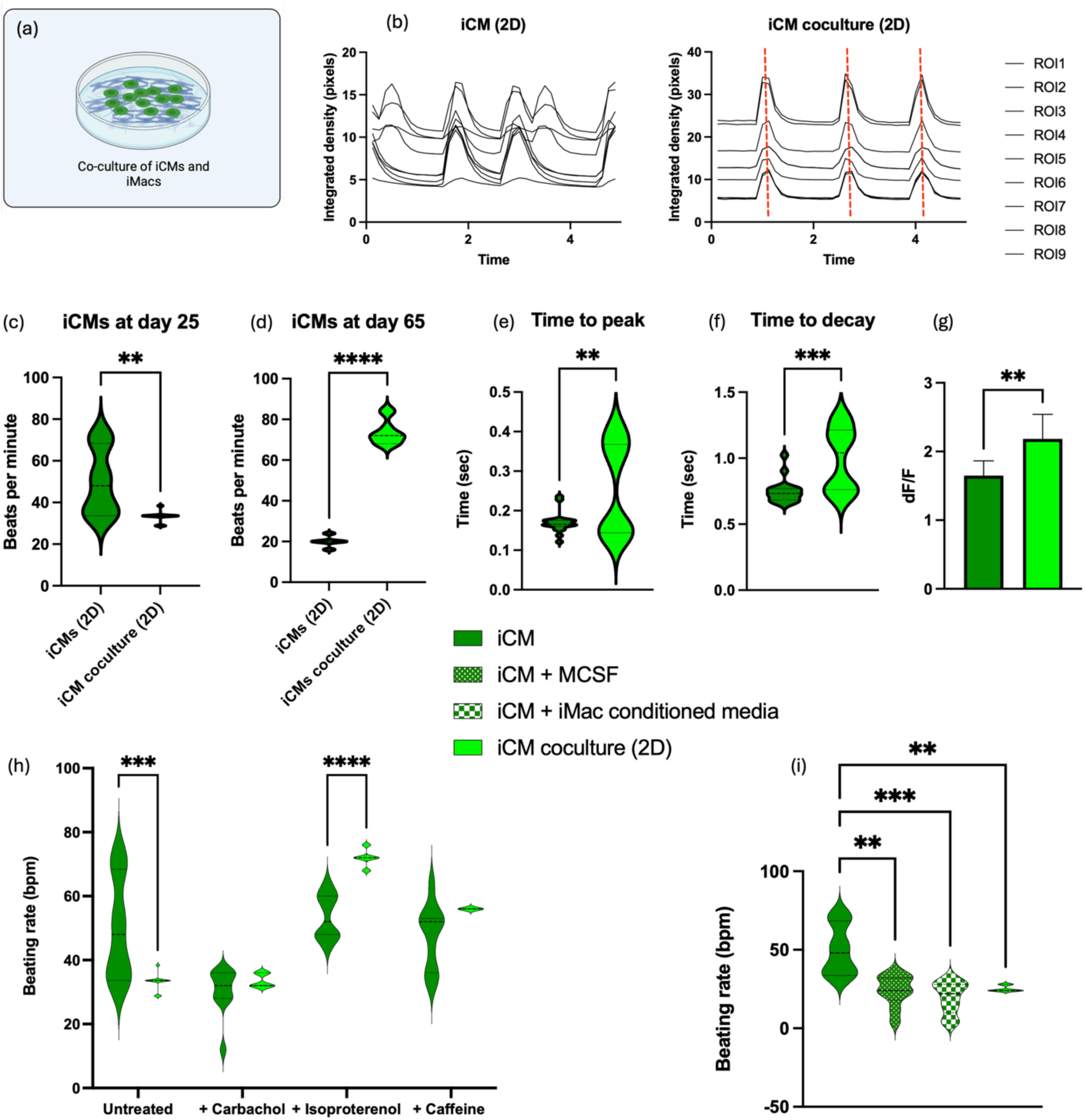
iMacs electrically couple with iCMs and facilitate synchronous beating in addition to improved calcium handling. (**a**) Schematic representation of the experimental setup, (**b**) Representative plots of the calcium transients in iCMs and iCMs in co-culture with iMacs. Nine regions of interest are assessed, and the peaks denote individual spontaneous beat of the cells. Red lines highlight the synchronous beating of iCMs in co-culture with iMacs, (**c**) Quantified beat rate of iCMs and iCMs in co-culture with iMacs at day 25, (**d**) Quantified beat rate of iCMs and iCMs in co-culture with iMacs at day 65, (**e**) Quantification of time to calcium peak, (**f**) Quantification of time to calcium decay, (**g**) Quantified amplitude of calcium activity (dF/F), (**h**) Quantified beat rate of iCMs and iCMs in co-culture with iMacs in response to treatment with carbachol, isoproterenol and caffeine. (**i**) Quantified beat rate of iCMs, iCMs cultured with M-CSF, iCMs cultured with iMacs conditioned media and iCMs in co-culture with iMacs. Statistical analysis - Student T test with Welch’s correction for c, d, e, f and g; Two-way ANOVA with Tukey’s multiple comparison for h; one-way ANOVA with Tukey’s comparison test for i. * represents P<0.05, ** represents P<0.01, *** represents P<0.001 and **** represents P<0.0001. iCMs and iMacs generated using SFCi55 iPSC line.

Following this initial observation, calcium transient read-outs were used to quantify the beat rate of iCMs and iCMs in co-culture with iMacs. Using the same approach as before, different regions of interest were chosen in individual wells of iCM cultures. On day 25 of differentiation, distinct patches of iCMs beat at vastly different paces, with an average of 49.92 ± 15.87 bpm (Fig. 5 (c)) (Supplementary video 2). However, iCMs in co-culture with iMacs were found to beat at an average of 33.12 ± 2.72 bpm. It is worth noting that iCMs in co-culture with iMacs exhibited strikingly low variability in terms of their rate of spontaneous beating (Fig. 5 (c)), indicating synchronous beating of various patches (Supplementary video 3). Furthermore, other calcium handling properties such as time to peak calcium transients and time to decay of calcium transients were both found to be more prolonged in iCMs that were in co-culture with iMacs (Fig. 5 (e and f)). Similarly, the intensity of calcium transients (dF/F) was also found to be greater in iCMs that were in co-culture with iMacs (Fig. 5 (g)).

Several compounds can be used to alter the beat rate of cardiomyocytes, and this effect can be induced by targeting different ion channels and processes. Subsequently, it was intriguing for us to investigate whether iCMs in co-culture with iMacs also responded to known drugs such as carbamylcholine (carbachol), isoproterenol and caffeine. In agreement with previous reports [66] [67, 68], both iCMs and iCM/iMacs co-cultures exhibited increased beat rates with carbachol treatment, and reduced beat rates with isoproterenol treatment (Fig. 5 (h)). Caffeine is also a commonly used drug that increases the beat rate of iCMs [69] and improves their active calcium transients [70]. In our study, although iCMs did not show a notable increase in beat rate on exposure to caffeine (47.6 ± 10.2 bpm), iCM/iMacs co-cultures were highly sensitive and displayed a steady beat rate of 56 ± 0 bpm (Fig. 5 (h)). Despite iCMs being known to exhibit poor calcium handling [71], our results suggest the addition of iMacs show great potential in improving the calcium handling of iCMs, with highly robust beat rates and very little variability. To further confirm that the synchronous beating of iCMs was facilitated through electrical coupling with iMacs and not just through paracrine factors, iCMs were also cultured for 7 days with M-CSF, and with iMacs conditioned media. Synchronous beating with very little variability could only be seen when iCMs were in co-culture with iMacs (Fig. 5 (i)) (Supplementary video 3), whereas iCMs cultured with macrophage colony stimulating factor (M-CSF) (Supplementary video 4) and iMacs conditioned media (Supplementary video 5) displayed high variability in beat rates (24 ± 10 bpm and 19.6 ± 10 bpm respectively) (Fig. 5 (i)).

Previous studies also show that prolonged culture time (>50 days) improved iCM maturity, facilitating elongation of cells [72] and causing reduced beat rate [73, 74]. In agreement to these findings, iCMs kept in culture for 65 days showed a highly reduced beat rate of 20 ± 2.6 bpm, with very little variability (Fig. 5 (d)). iCMs in co-culture with iMacs displayed a beat rate of 73.6 ± 6 bpm (Fig. 5 (d)), within the physiological range of beating for adult human hearts [75]. Taken together, these findings strongly emphasise the functional maturity displayed by iCMs in co-culture with iMacs, with highly synchronous beating, robust response to drug treatments and physiologically accurate beat rates.

### iMacs remain highly quiescent, with notable functional activation when electrically stimulated in co-culture with iCMs

Owing to the highly immature phenotype of iCMs, their use in translational research is debated, with iCMs being functionally closer to foetal rather than adult cardiomyocytes [76]. Although the addition of iMacs showed some structural and functional improvements in iCMs, next assessed combinatorial effects of iMacs addition and electrical stimulation (ES) in improving both macrophage and cardiomyocyte function. Out of a multitude of approaches used to improve iCM maturation, in this study the effect of periodic and continuous electrical stimulations was explored (Supplementary Figure 6 (a)). Similar to the characteristics of native myocardium [77, 78], biphasic pulsatile waveforms of 2 ms pulse width, ± 2.5 V and 2 Hz frequency were used. Hence, for periodic stimulation, iCMs were exposed to ES for 1 hour for 5 days (iCM_PR1) and for continuous stimulation, iCMs were exposed to ES for 5 full days (iCM_PR2) (Supplementary Figure 6). On assessing the gene expression profiles, iCMs undergoing continuous stimulation, iCM_PR2, were found to express more *TNNT2* (95.4% higher), *MYH6* (89.4% higher), *MYH7* (97.5% higher) and *SERCA2* (97% higher), in comparison to iCMs (Supplementary Figure 6 (b and c)). Additionally, the *MYH7*/*MYH6* ratio was 62.7% upregulated in iCM_PR2 compared to iCMs (Supplementary Figure 6 (c)), indicating a high upregulation of key genes related to iCM maturation, when electrically stimulated continuously for 5 days.

Given that iMacs in co-culture with iCMs would also experience similar ES, it was pertinent to assess iMacs individually under ES (iMacs_5d_ES). Notably, upon 5 days of ES, iMacs expressed 82.7% more *TGFB1*, 84.5% more *MRC1* and 79.6% more *IL10*, in comparison to unstimulated iMacs, displaying a highly pro-regenerative response (Fig. 6 (a)). The anti-inflammatory response was also highlighted by the downregulation in *NOS2* (92%, compared to iMacs) and no significant difference in *TNF* gene expression (Fig. 6 (d)). Additionally, genes associated with functional activation of iMacs were also assessed. *MMP9* was 91% lower in iMacs_5d_ES (Fig. 6 (c)), indicating homeostasis and a potential for robust matrix production [35]. Furthermore, iMacs_5d ES also exhibited 83.3% upregulation of *CCR2*, 36% upregulation of *TRPV4* and 72.8% upregulation of *GJA1* (Fig. 6 (c)), demonstrating potential activation of gap junctions. These findings are particularly interesting in the context of the myocardium, as both the CCR2^+^ and CCR2^-^ macrophages express TRPV4 and GJA1 in abundance [13, 44]. No significant difference was noted in the expressions of *VEGFA* and *AREG*, between iMacs_5d_ES and untreated iMacs (Fig. 6 (c)). Overall, this study is first of its kind, examining the effects of providing iMacs with cardiac-specific ES. These results not only reiterate the highly anti-inflammatory phenotype of iMacs under ES, but also underline the safety of delivering ES to iMacs.

**Fig. 6.**
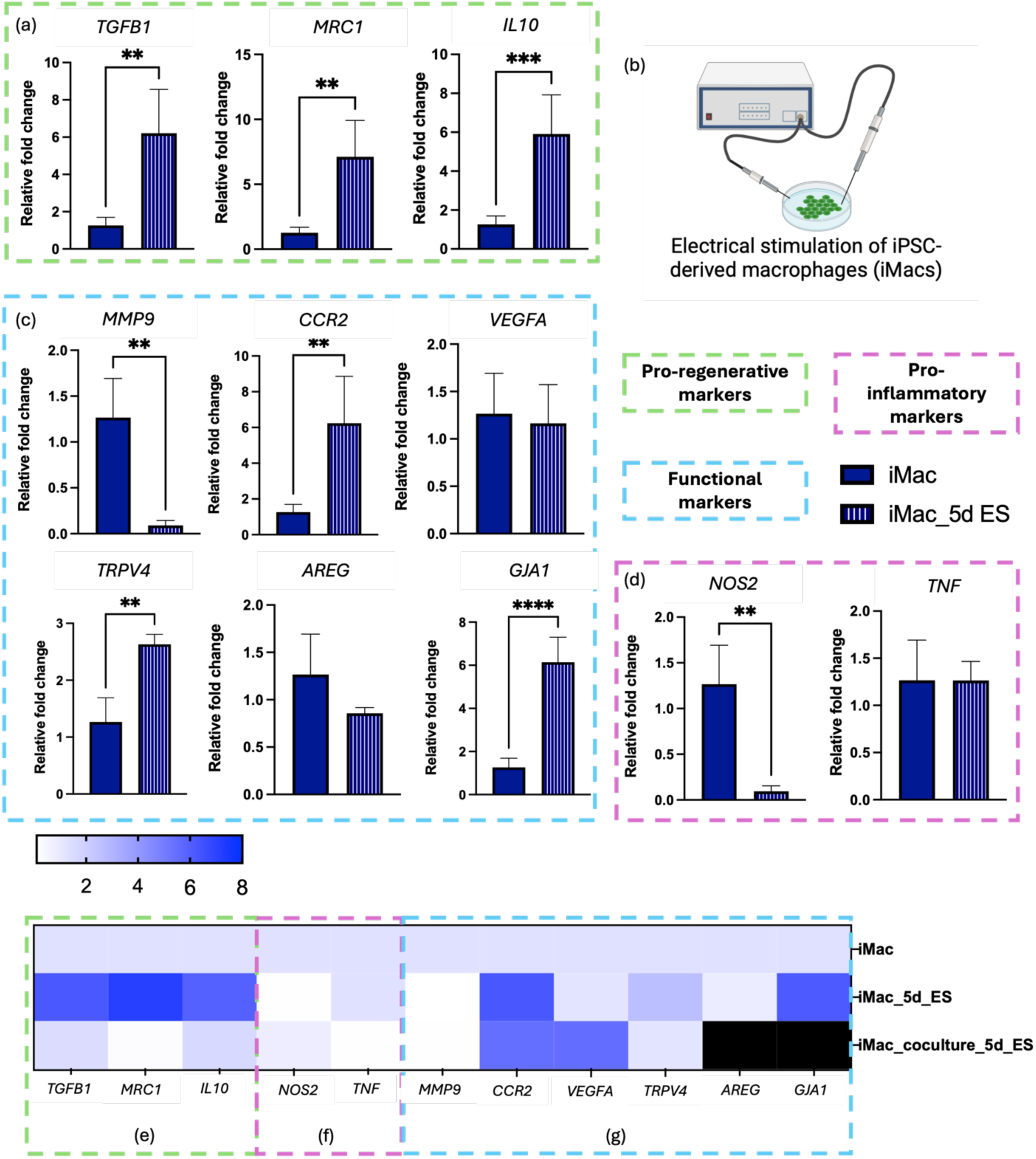
iMacs are highly pro-regenerative when exposed to 5 days of electrical stimulation. **(a)** Expression of a set of pro-regenerative genes, quantified using qPCR (**b**) Schematic representation of the experimental set up, (**c**) Expression of a set of macrophage functional genes, quantified using qPCR (**d**) Expression of a set of pro-inflammatory genes, quantified using qPCR, Heat map comparing the expressions of a set of (**e**) pro-regenerative, (**f**) pro-inflammatory and (**g**) functional genes between iMacs, iMacs under 5 days of ES and iMacs in co-culture with iCMs under 5 days of ES. Black boxes indicate relative mRNA expression over 40-fold. Statistical analysis - Student T test with Welch’s correction. ** represents P<0.01, *** represents P<0.001 and **** represents P<0.0001.

After individually characterising electrically stimulated iCMs and iMacs, co-cultures of iCMs and iMacs were subjected to 5 days of ES. The first set of assessment was performed on iMacs in co-culture with iCMs, to evaluate the phenotype of iMacs. While continuous ES for 5 days caused an upregulation in the expression of pro-regenerative genes (Fig. 6 (e)), iMacs in co-culture with iCMs exposed to the same ES were highly quiescent, with no significant pro-regenerative responses (Fig. 6 (e)). The quiescence of iMacs could also be seen in their expression of pro-inflammatory genes such as *NOS2* and *TNF* (91.6% downregulation in comparison to iMacs) (Fig. 6 (f)). Regardless, strong functional activation was also noted in electrically stimulated iMacs_co-culture, with a 77% upregulation in *VEGFA* expression, 97.5 % upregulation in *AREG* expression, and a 92.8% upregulation in *GJA1*, in comparison to untreated iMacs (Fig. 6 (g)), all genes of critical importance in CRMs [13, 44, 46, 79]. It is worth noting that, phenotypically and functionally, iMacs and electrically stimulated iMacs_co-culture were similar, except for the highly upregulated expressions of *VEGFA* and *AREG* in stimulated iMacs_co-culture. Moreover, on examining electrically stimulated iMacs for their expression of CRM-specific and cardiac developmental genes, the expressions were either lower than unstimulated iMacs_co-culture, or not significantly different from unstimulated iMacs_co-culture (Supplementary Figure 7). Hence, electrically stimulated iMacs_co-culture were not examined for their expression of cardiac developmental and CRM-specific genes.

Furthermore, the secretome from iMacs in co-culture with iCMs, with and without ES were collected and assessed for the presence of pro-inflammatory cytokines. While negligible amount of TNFα was found in the secretome of both the test conditions, a 94.1% downregulation of IL6 and 97.1% downregulation of IL10 was found in electrically stimulated iMacs_co-culture compared to unstimulated co-cultures (Supplementary Figure 8). Nonetheless, in this study, no inflammatory response was found, which in part can be associated with the synergistic effect of iMacs being in physical contact with iCMs, as well as the potential anti-inflammatory factors released by iCMs. Taken together, these findings further highlight the phenotypic quiescence of electrically stimulated iMacs in co-culture with iCMs, with remarkable functional activation, exhibiting high *VEGFA, AREG* and *GJA1* expressions. These findings underline the benefits of ES in conjunction with iCM culture for functional improvement of iMacs.

### Co-culture with iMacs yields similar level of maturation in iCMs as 5 days of electrical stimulation

After seeing encouraging functional activation in electrically stimulated iMacs_co-culture, we next focused on analysing the maturation of electrically stimulated iCMs in co-culture with iMacs (Fig.7 (a)). After 5 days of ES, iCMs and iCM in co-culture with iMacs showed improved beat rates, with very little variability (Fig. 7 (b))). Notably, paced iCMs beat at 80 ± 22 bpm compared to paced co-cultures 86 ± 5 bpm (Fig. 7 (b)). Next, the expressions of cardiomyocyte maturation genes were assessed. As previously described (Supplementary Figure 6), 5 days of ES resulted in a notable increase in the iCM maturation genes evaluated. In comparison, paced iCMs in co-culture exhibited comparable levels of *TNNT2* (94.5% more than unstimulated iCMs) (Fig. 7 (c)). Similarly, the ratio of *MYH7*/*MYH6* was also comparable between paced iCMs and paced iCMs in co-culture, with 71.7% upregulation compared to iCMs (Fig. 7 (c)). However, the expression of *SERCA2* was 79.6% higher in paced iCMs than paced iCMs in co-culture (Fig. 7 (c)). *SERCA2* expression was comparable between electrically stimulated iCMs in co-culture and unstimulated iCMs in co-culture (Fig. 7 (c)). Electrically stimulated iCMs and co-cultures exhibited a similar aspect ratio as unstimulated iCMs in co-culture with iMacs (Fig. 7 (d)), indicating robust cell elongation. On assessing the morphology of iCMs, the aspect ratio increased by 52.6% upon ES (increase from 1.18 ± 0.24 to 2.39 ± 0.46) (Fig. 7 (d)). However, iCMs in co-culture with iMacs did not show statistically significant upregulation in the aspect ratio upon ES (2.86 ± 0.61 for unstimulated co-cultures; 3.15 ± 0.75 for stimulated co-cultures) (Fig. 7 (d)). Furthermore, while iMacs addition showed improvement in the intensity of calcium transients in the absence of ES (Fig. 7 (e)), under ES, there was no significant difference in the calcium intensity between iCMs and iCMs in co-culture with iMacs (Fig. 7 (e)).

**Fig. 7.**
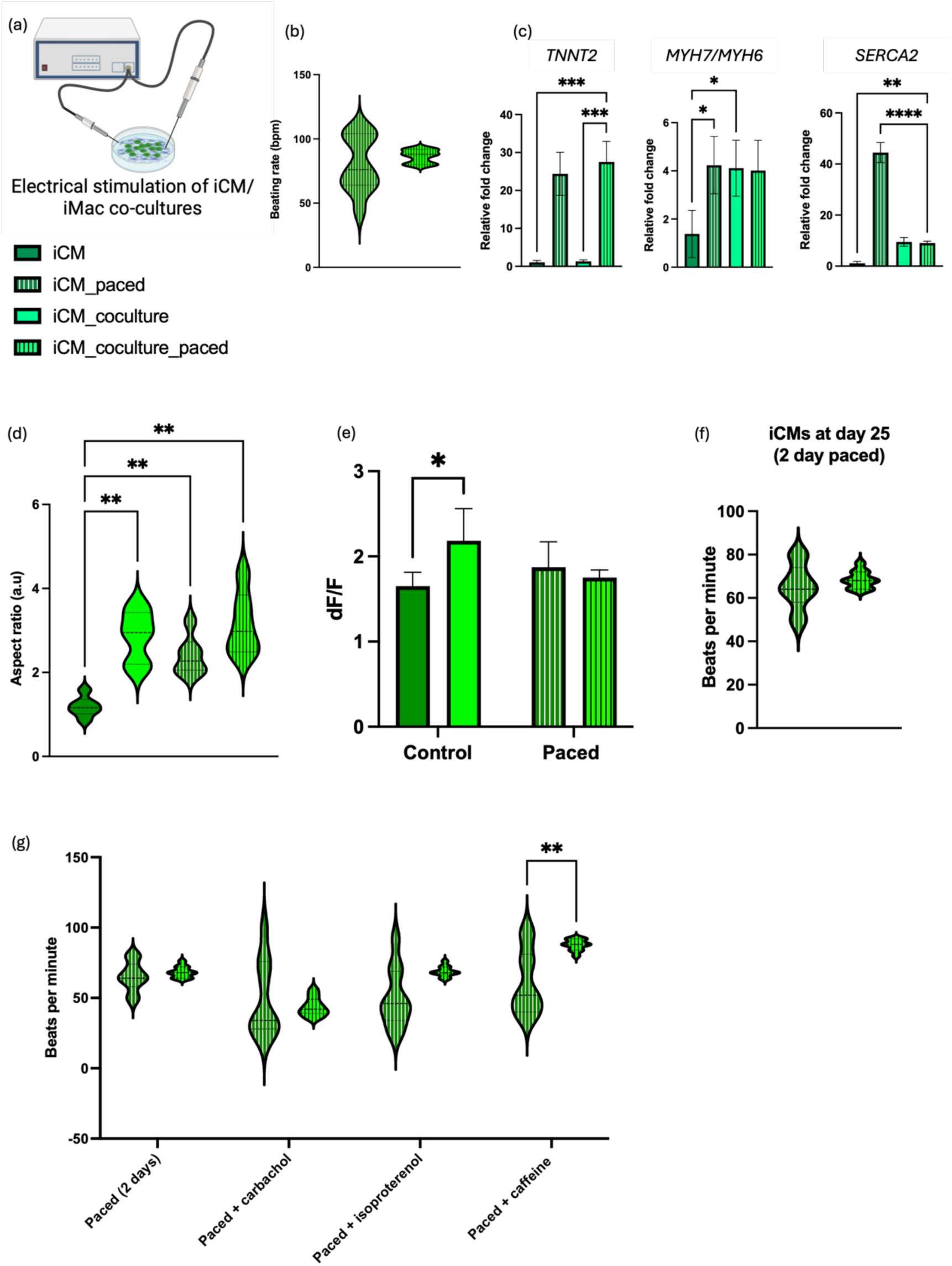
Co-culture with iMacs yields similar level of maturation in iCMs as 5 days of electrical stimulation. (**a**) Schematic representation of the experimental set up, (**b**) Quantified beat rate of iCMs and co-cultures after 5 days of ES, (**c**) Expression of a set of cardiomyocyte maturation genes, using qPCR (**d**) Aspect ratio of iCMs and co-cultures, with and without 5 days of ES, (**e**) Intensity (dF/F) of calcium signalling by iCMs and co-cultures at day 25 (control) and after 5 days of ES (paced), (**f)** Quantified beat rate of iCMs and co-cultures after 2 days of ES, (**g**) Quantified beat rate of iCMs and co-cultures in response to treatment with carbachol, isoproterenol and caffeine, after 2 days of ES. Statistical analysis - Students T test with Welch’s correction for b and f; One-Way ANOVA with Tukey’s multiple comparison for c and d; Two-way ANOVA with Tukey’s multiple comparison for e and g. * represents P<0.05, ** represents P<0.01, *** represents P<0.001 and **** represents P<0.0001.

Continuous ES evidently yielded positive responses in iCM maturation gene expression, morphology (elongation) and beat rate. However, this led us to question whether iCM/iMacs co-cultures require the entire 5 days of ES to exhibit robust calcium handling responses. Hence, iCM/iMacs co-cultures were electrically stimulated for only 2 days and the beat rate and calcium handling functions of iCMs were evaluated. After 2 days of ES, iCMs displayed a beat rate of 65 ± 10 bpm while paced iCM/iMacs co-cultures displayed a robust 68 ± 2 bpm, with very little variability, reiterating the exceptional ability of iMacs to electrically couple with iCMs and facilitate synchronous beating (Fig. 7 (f)). Furthermore, we also explored the response of electrically stimulated (for 2 days) iCMs to the exposure of known drugs. On treating electrically stimulated iCMs and iCM/iMacs co-cultures with carbachol, the beat rate was found to be 48 ± 27 bpm for iCMs and 43 ± 6 bpm for iCMs in co-culture with iMacs (Fig. 7 (g)). Next, isoproterenol treatment increased the beat rate to 50 ± 21 bpm for iCMs and 68 ± 3 bpm for co-cultures (Fig. 7 (g)). Furthermore, caffeine treatment was found to increase the beat rate to 61 ± 22 bpm for iCMs and 87 ± 3 bpm for co-cultures, yielding a 30.1% increase in co-cultures (Fig. 7 (g)). Collectively, 2 days of ES was found to yield robust responses in iCMs in co-culture with iMacs, with notably low variability on treatment with carbachol, isoproterenol and caffeine, while iCMs showed high variability in the beat rate (Fig. 7 (g)).

These observations emphasise that while iCMs remain fairly immature even with 2 days of ES, iCMs in co-culture with iMacs show improved maturation, comparable to the maturation achieved after 5 days of ES, in terms of their gene expression profile and elongated morphology. Moreover, with 2 days of ES, iCMs in co-culture with iMacs also carry out robust calcium handling functions. Taken together, these results highlight iMacs as an excellent support cell type to improve iCM maturation and function, by enhancing synchronous beating, increasing structural maturation (by facilitating elongation), improving calcium handling, and upregulating the expression of maturation genes.

### ECTs containing iMacs display improved iCM elongation, beat rate and tissue compaction

Having observed the positive impact of co-culturing iMacs and iCMs, we then fabricated engineered cardiac tissues (ECTs) using collagen and Matrigel^®^ containing iMacs and iCMs, to evaluate interactions in a 3D setting. Before making the ECTs, we confirmed that the collagen and Matrigel^®^ hydrogel combination was not triggering an inflammatory response in iMacs (Supplementary Figure 9). By performing flow cytometry, we observed that pro-inflammatory markers such as CD80 and CD86, as well as anti-inflammatory markers such as CD163 and CD206 were similar between iMacs that were previously encapsulated in the collagen and Matrigel^®^ hydrogel and control iMacs that were cultured on tissue culture plastic (Supplementary Figure 9). Subsequently, using isogenic iCM and iMacs sources (derived from SCFi55 iPSCs), two groups of ECTs were fabricated – one containing only iCMs and the other containing both iCMs and iMacs (Fig. 8 (a, c and d)). While both iCM and iCM/iMacs ECTs showed alignment to the axis of anchorage, more cells (greater intensity) displayed preferential alignment in iCM/iMacs ECTs (Fig. 8 (b)). On assessing the elongation of iCMs in the ECTs, iCMs were found to be more elongated in the iCM/iMacs ECTs, with a 36.7% increase in the aspect ratio (Fig. 8 (e)). However, in comparison to iCMs in monolayer, iCMs in ECTs displayed significantly greater elongation (aspect ratio of 3.18), highlighting structural maturation. Similar elongation of iCMs in ECTs has been shown in previous work, although not quantified [9, 11]. Both iCM ECTs and iCM/iMacs ECTs displayed physiologically relevant beat rates (Fig. 8 (f)); range highlighted in light green) that were comparable to each other. While iCM ECTs beat at 68 ± 10 bpm, iCM/iMacs ECTs beat at 72 ± 4 bpm, distinctly demonstrating improved synchrony and reduced variability in the beat rate. Unsurprisingly, the reported beat rates for ECTs vary vastly, owing to the large degree of variability across iPSC lines. While Hansen *et al.* reported a range of beat rates from 20 to 162 bpm [9], other reports include 42 bpm [11], 50 bpm [80] and 49 bpm [81]. However, Mannhardt *et al*. reported a baseline beat rate of 61 ± 2 bpm (close to the value reported in this study), for fibrin-based ECTs [82]. Finally, the degree of compaction of the tissues, which is a hallmark of tissue remodeling, matrix deposition and contractile performance [83–85], was also assessed based on the cross-sectional area of the tissue. As early as day 3 of the culture, ECTs containing iMacs showed improved compaction, which was found to be further accelerated by day 7 (Fig. 8 (g)). Notably, iCM ECTs did not compact over the period of 7 days (Fig. 8 (g)). With more recent studies highlighting the advantages of adding macrophages (iMacs [86] and ESC-derived macrophages [23]), these results illuminate the benefit of adding iMacs to ECTs, to improve the beat rate and compaction of the tissues, in addition to facilitating iCM elongation.

**Fig. 8.**
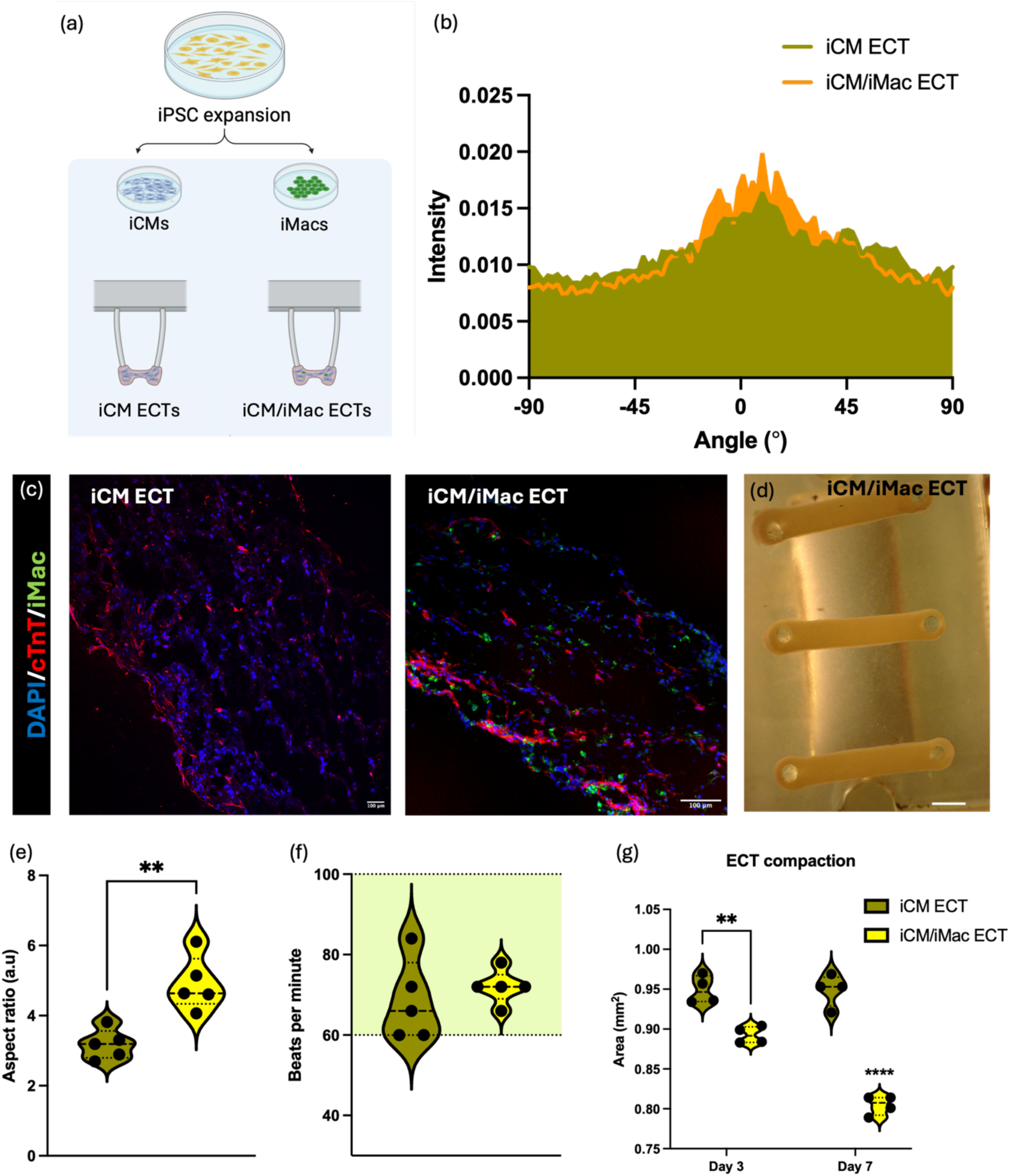
Addition of iMacs into engineered cardiac tissues improve cell alignment, beat rate and compaction of the tissues. (**a**) Schematic representation of the experimental set up, (**b**) Quantification of the alignment of iCM and iCM/iMacs ECTs. Green area indicates the alignment of iCM ECTs; orange area indicates the alignment of iCM/iMacs ECTs, (**c**) Representative images of iCM and iCM/iMacs ECTs stained with cardiac troponin T. iMacs differentiated from SFCi55-ZsGr iPSC line. Scale bar – 100 µm, (**d**) Representative images of iCM/iMacs ECTs in culture. Scale bar – 1 mm, (**e**) Quantified aspect ratio of iCMs in iCM ECTs and in iCM/iMacs ECTs, (**f)** Quantified beat rate (in bpm) of iCM and iCM/iMacs ECTs. Light green area highlights range of human heartbeat rate, (**g**) Compaction of ECTs over a period of 7 days. Statistical analysis - Students T test with Welch’s correction for e and f; Two-way ANOVA with Tukey’s multiple comparison for g. ** represents P<0.01 and **** represents P<0.0001.

## CONCLUSION

The field of cardiac immunology has gained more interest in the recent past, owing to the discovery of key interactions between cardiomyocytes and macrophages, and the role of macrophages (in particular, CRMs) in maintaining homeostasis within the myocardium. Most of the reported functions of CRMs are using murine models, with very little known about CRMs in humans. To deepen our understanding of human CRM biology, *in vitro* cultures can be used as an effective tool. The primary objective of this study was to understand how cardiomyocytes and macrophages interact with each other *in vitro*, using iCMs and iMacs from the same iPSC parent line (SFCi55). To efficiently simulate a cardiac environment in a 2D setting *in vitro*, two approaches were chosen - providing iMacs with iCM conditioned media (iMacs_CM) and performing co-cultures of iMacs and iCMs (iMacs_co-culture). In this comprehensive exploration, the most profound impact in terms of iMac maturation and function was seen when iMacs and iCMs were in co-culture, showing that the effects of CRMs in the heart extend beyond paracrine signalling. Despite high cell densities of iCMs, iMacs integrated well into iCM cultures, particularly into beating patches. The expression of Cx43 in both cell types further emphasised potential communication through gap junctions, crucial for myocardial function. It should be emphasised that, iMacs_co-culture not only displayed a reduced level of pro-inflammatory genes (*TNF* and *NOS2*) compared to control iMacs, but also expressed a robust increase in expression of CRM-specific genes (*TIMD4*, *LYVE1* and *FOLR2*) and cardiac developmental genes (*GATA4*, *TBX18* and *TBX5*), indicating overall quiescence, a key feature of TRMs.

In addition to characterising the response of iMacs, the phenotypic and functional changes in iCMs was also investigated. In co-culture with iMacs, iCMs displayed an elongated morphology, in addition to an increase in the expression of maturation markers such as *SERCA2*, the ratio of *MHY7*/*MHY6* and improved calcium handling. Furthermore, one of the most significant drawbacks of iCM cultures - asynchronous beating of the cells - was mitigated upon co-culture with iMacs, wherein synchronous beating of iCMs could be seen, owing to electrical coupling between iMacs and iCMs. To that end, the gene and protein expression profiles of both cell types gave good indicators of their maturation and cardiac resident function. Subsequently, we sought to address the limitation of poor iCM maturity in *in vitro* cultures [76]. To achieve improvement in iCM maturity, electrical stimulation (ES) was chosen, as over 2 decades of research backs the efficacy of this approach [78]. To maintain the same systematic approach, the effect of ES on iCMs, iMacs and iCM/iMacs co-cultures was investigated, by adopting a physiologically accurate ES regimen resembling the characteristics of native myocardium. The ES regimen that yielded the greatest improvement in the expression of iCM maturation genes also drove a highly pro-regenerative response in iMacs. Moreover, functional activation markers, including *MMP9, CCR2, TRPV4,* and *GJA1*, were notably upregulated, demonstrating the multifaceted impact of ES on iMacs. Furthermore, the examination of iMacs from electrically stimulated co-cultures revealed that they maintained quiescence, in addition to expressing pro-inflammatory genes at significantly reduced levels. Electrically stimulated iMacs in co-culture with iCMs also demonstrated exceptional functional activation, emphasising high *VEGFA*, *AREG*, and *GJA1* expressions. Interestingly, functional readouts of iCMs, such as calcium handling (dF/F), elongation and maturation gene expressions were found to be comparable between electrically stimulated iCMs and iCMs in co-culture with iMacs, further highlighting the symbiosis between iMacs and iCMs (Fig. 7(c, d, e)). These findings not only reiterated the anti-inflammatory phenotype of iMacs under ES but also highlighted the safety of delivering ES to iMacs, reinforcing the benefits of ES in conjunction with iCM culture for functional improvement of iMacs.

Having seen excellent interactions between iCMs and iMacs in a 2D setting, our next focus was to study the effect of iMacs addition on ECTs. Although majority of conventional ECT models only focused on cardiomyocytes, or a combination of cardiomyocytes and cardiac fibroblasts [58, 87, 88], newer studies underline the immense benefits of adding iMacs into ECTs in terms of improving tissue function [23] and vascularisation [89]. In line with these findings, addition of iMacs significantly reduced the variability in the beat rate of the tissues, improved cell alignment and iCM elongation, as well as remarkably increased tissue compaction. These results indicate that in comparison to conventional ECTs, ECTs with iMacs are be more enhanced and physiologically relevant.

In conclusion, the findings of this work show that iMacs in co-culture with iCMs exhibit a more CRM-like phenotype and iCMs display improved maturation and calcium handling in this co-culture. Additionally, these findings also highlight iMacs as excellent support cells in ECTs, contributing to improved tissue function and providing a foundation for advancing cardiac immunoengineering. The findings of this study open up new avenues for further research in the field of cardiac tissue engineering and regenerative medicine to help understand the complex interactions between macrophages and other non-myocytes, to develop true multicellular healthy and injured cardiac models.

## MATERIALS AND METHODS

### Differentiation of iPSCs to macrophages (iMacs)

All experiments were performed using iPSC lines SCFi55 (accession: CVCL_UL47) and SFCi55-ZsGr (assession: CVCL_UL48; tagged with Zeiss green) [90]. iPSCs were expanded using mTESR™ media (Stemcell Technologies) in Matrigel^®^ (Corning) coated plates, with the addition of 10 µM ROCK inhibitor (Stemcell Technologies) on days of passaging. For embryoid body (EB) formation, one well of 80-90% confluent iPSCs (at P18 to P22) were lifted and plated on an ultra-low attachment (ULA) plate (Greiner Bio-One). On day 0, cells were cultured in mTESR™ media supplemented with BMP-4 (50 ng/ml), VEGF (50 ng/ml) and SCF (20 ng/ml). On day 2 of differentiation, an additional 0.5 ml media was added per well, with concentrations of BMP-4 (Biotechne), VEGF (Biotechne) and SCF (Biotechne) adjusted to make up for the entire media volume. On day 4 of differentiation, the EBs were collected and plated on 0.1 % Gelatin (Stemcell Technologies) coated 6-well plates at a concentration of 12-15 EBs per well, to initiate haematopoiesis induction. The plated EBs were cultured in X-VIVO media (Lonza) supplemented with M-CSF (100 ng/ml; Miltenyi), IL3 (25 ng/ml; Biotechne), Glutamax (2 mM; Gibco), 2% penicillin/streptomycin (Gibco) and beta mercaptoethanol (55 mM; Gibco). At about 14 days after the initiation of haematopoiesis, the non-adherent cells or macrophage precursors (pre-macs) present in the wells were collected and plated at a density of 500,000 cells per well of a 6-well plate and cultured in X-VIVO media, supplemented with 100 ng/ml M-CSF, 1% Glutamax and 2% penicillin/streptomycin for 7 days, to obtain macrophages. iMacs were predominantly differentiated from SFCi55-ZsGr iPSC line (tagged with Zeiss green).

### iMacs treated with iCM conditioned media

For regular iMacs culture, pre-macs (non-adherent cells released by EBs from day 14 of the initiation of haematopoiesis) were collected and plated at a density of 500,000 cells per well of a 6-well plate and cultured in X-VIVO media, supplemented with 100 ng/ml M-CSF, 1% Glutamax and 2% penicillin/streptomycin (iMacs media) for 7 days. Conditioned media was collected from iCMs between day 20 and 30 of culture and centrifuged at 650 G, to remove dead cells and debris, before storing at -80 °C. For iCM conditioned media studies, pre-macs were collected and plated at a density of 500,000 cells per well of a 6-well plate or 100,000 cells per well of a 24-well plate and cultured with 1:1 ratio of iMacs media and iCM conditioned media supplemented with 100 ng/ml M-CSF. Media change was performed at day 3 of the culture and iMacs_CM were used for experiments at day 7 of culture. For immunofluorescence imaging with iCM conditioned media treatment, pre-macs were plated at a density of 500,000 cells per well of a 35mm µ-dish (IBIDI^®^) and cultured with 1:1 ratio of iMacs media and iCM conditioned media supplemented with 100 ng/ml M-CSF. BDMs or untreated iMacs were used as controls and were cultured at the same cell density as iMacs_CM. BDMs were cultured in RPMI media with 50 ng/ml M-CSF, 10% FBS and 2% pen/strep (BDM media).

### Differentiation of iPSCs to cardiomyocytes (iCMs)

SFCi55 iPSCs were used to derive iCMs, using the GiWi protocol [91]. Briefly, iPSCs were expanded until 95-100% confluency. Mesodermal specification was induced using 10 µM CHIR99021 (Stem Cell Technologies). On day 0 of differentiation, RPMI 1640 (Gibco) media containing 2% B-27 minus insulin (Gibco) (RPMI-i) and 10 µM CHIR99021 was added to the iPSCs. CHIR99021 containing media was removed from the cultures after 24 hours and replaced with RPMI-i. On day 3 of the differentiation, RPMI-i was supplemented with 5 mM IWP2 and was added to the cultures, to initiate cardiac mesoderm induction. Another media change was performed on day 5, with RPMI-i and finally, on day 7, cells were treated with RPMI media with complete B-27 (RPMI+i). Media was changed every 3 days, and the cells were used for experiments between days 21-30 of differentiation.

### Co-culture of iMacs and iCMs

For performing co-cultures of iMacs and iCMs, iCMs were differentiated and cultured for up to 20 days before replating them at a density of 200,000 cells/cm^2^, on Matrigel^®^ coated plates. Replated iCMs were allowed to attach for 48 hours before adding pre-macs to the cultures, at a 5:1 ratio of iCMs to iMacs. Co-cultures of iCMs and iMacs were cultured in RPMI+i (RPMI media with 2% B27™ and 2% pen/strep) supplemented with 100 ng/ml M-CSF. Cells were kept in culture for 7 days before performing experiments and a complete media change was performed at day 3. On day 7 of co-culture, cells were lifted from the plates by incubating them with 0.25% Trypsin-EDTA solution (Thermo Fisher Scientific) at 37 °C for 9 minutes. The lifted cell suspension was centrifuged at 300 G for 5 minutes, followed by resuspension in FACS sorting buffer (Ca/Mg^2+^ free PBS with 2.5 mM EDTA, 25 mM HEPES, 1% heat inactivated FBS and 1% pen/strep). Since iMacs were generated using SFCi55-ZsGr iPSCs and iCMs were generated using SFCi55 iPSCs, the resuspended cells were FACS sorted. Positive sorting was done for cells expressing Zeiss green (iMacs) and the flow through cells collected as iCMs. FACS sorted iMacs and iCMs were lysed using Trizol™, to perform RT-qPCR.

### Polarisation of iMacs

Polarisation was induced in iMacs by treating them with certain factors that trigger a pro-inflammatory and anti-inflammatory response. To drive a pro-inflammatory response, iMacs were treated with 100 ng/ml lipopolysaccharide (LPS) and 50 ng/ml of interferon gamma (IFNγ) (Biotechne) for 24 hours. Similarly, to drive an anti-inflammatory response, iMacs were treated with 20 ng/ml IL4 (Biotechne) for 24 hours, with untreated iMacs used as control. The polarisation potential of iMacs was evaluated through ELISA and flow cytometry. ELISA was performed to quantify the presence of cytokines such as TNFα, IL6 and IL10 in the secretome, and flow cytometry was performed to quantify the expression of CD80, CD86, CD163 and CD206.

### Phagocytic potential of iMacs

Pre-macs were plated on 24 well plates at a density of 300,000 cells/well and cultured for 7 days in the presence of 100 ng/ml M-SCF. Red zymosan particles (Phagocytosis Assay Kit, (Red Zymosan) Abcam) were added to the culture and incubated for 3 hours to facilitate phagocytosis. After incubation, the cells were washed with ice cold phagocytosis assay buffer and retrieved from the plates using a cell scraper. Flow cytometry was used to quantify the population of cells positive for the red zymosan particles (550 nm). BDMs were used as control.

### Electrical Stimulation bioreactor setup

The bioreactor set up to generate ES was custom-built for previous studies in the lab [92]. Briefly, a 1 mm hole was drilled one end of each of the twelve 22 mm long cylindrical carbon electrode, to secure a platinum-iridium wire. The platinum wires and carbon electrodes were kept in place by immobilising them using PDMS Sylgard^TM^ 184, using a 3D printed mold. A lid adaptor that fits into the lid of a 6-well plate was 3D printed. The lid adaptor allowed the attachment of platinum-iridium wires and carbon electrodes, through PDMS fillers. The carbon rod electrodes were kept at 15 mm distance between each other and facilitated the generation of ES to a 6-well plate. The lid adaptor attached to the lid of the 6-well plate through six screws, and had spaces in between the electrodes for optical accessibility with both upright and inverted microscopes. A 5 mm gap between the adaptor and the lid was accounted for holding the wiring. The source ground and positive wires connected externally to a bread board where a waveform generator (RSDG2000X, Radionics) and an oscilloscope (RSDS 1052 DL +, Radionics) were also connected. The optimised waveform used to improve iCM maturation was a pulsatile biphasic waveform with a voltage of ±2.5 V, 2 ms pulse width and 2 Hz frequency. To perform live imaging of samples, a plate stand was 3D printed to mount the 6-well plate and a Dino-lite digital microscope (RS Pro) was used.

### Collagen/Matrigel® hydrogel fabrication

The hydrogel combination used in this study was a mixture of bovine Type I collagen and Matrigel^®^. The hydrogels were fabricated using previously published methods [8, 93], to obtain a final solution containing 2 mg/ml concentration of bovine Type I collagen (Telocol^®^-6, Advanced Biomatrix) and 0.9 mg/ml Matrigel^®^ (Corning). Other components in the hydrogel mixture include 10x Minimum Essential Media (MEM), 10x PBS, sterile water, and the cell/media mixture. Once successfully neutralised, the hydrogel was allowed to self-assemble at 37°C for 2 hours.

### Engineered cardiac tissue bioreactor fabrication

The ECT bioreactor was fabricated by modifying a previously established model [8, 93]. First, this model was scaled down to fit into a well of a 6-well plate, for the bioreactor to integrate with the ES set up. Different iterations of designs were trialled by 3D printing using a 3D printer (Formlabs 2), until the optimal dimensions and design was achieved. The optimal design comprised of a two-part mold (Supplementary Figure 10 (a 1, 2)) which attached to each other (Supplementary Figure 10 (b)). Sylgard 184 Elastomer (PDMS; VWR International) was casted onto this mold and allowed to cure at 80°C for 1 hour, to obtain the ECT bioreactor with posts (Supplementary Figure 10 (c)). A frame (Supplementary Figure 10 (a 3)) was also developed to allow the attachment of the ECT device, for structural support (Supplementary Figure 10 (d)). Finally, the hydrogel mixture was casted on to the three cylindrical slots of the baseplate (Supplementary Figure 10 (a 4)) and the ECT device with the frame was attached to the baseplate (Supplementary Figure 10 (e)) to allow the hydrogel to form around the posts of the ECT bioreactor. The molds on which PDMS was casted (Supplementary Figure 10 (a 1 and 2) were machined from aluminium, whereas the frame (Supplementary Figure 10 (a 3)) and the baseplate (Supplementary Figure 10 (a 4)) were machined from polysulfone and polytetrafluoroethylene (PTFE), respectively. All the parts required for the ECT bioreactor (Supplementary Figure 10 (e)) were thoroughly cleaned and sterilised before cell culture experiments by treating them with 70% ethanol for 20 minutes, followed by 20 minutes of UV irradiation on either side of the parts. Tissues were fabricated using iCMs and iMacs, with a total cell number of 1 million per tissue.

### Isolation of PBMCs from buffy coats

The plasma depleted fraction of anti-coagulated blood (also known as buffy coats), from healthy human donors were obtained through Irish Blood Transfusion Board (IBTS), St. James’s Hospital, Dublin. The procurement of buffy coats was supported by a Level 2 Ethics approval from Trinity College Dublin, whereby, when required, the age and sex of the blood donors can be disclosed by IBTS.

Buffy coats were diluted at a 1:1 ratio using PBS and centrifuged at 1250 G for 10 minutes at room temperature, with the centrifuge de-acceleration occurring with brake off. The leukocyte containing layer of the buffy coat was extracted using a Pasteur pipette and diluted by half in sterile PBS. The diluted leukocytes were split across multiple 50 ml conical falcon tubes and layered over Lymphoprep^TM^ such that every falcon tube held 20 ml Lymphoprep^TM^ and 25 ml diluted leukocytes. Next, leukopcyte-Lymphoprep^TM^ mix was centrifuged at 800 G for 20 minutes with the brake off and the PBMC layer was carefully removed using a Pasteur pipette. The PBMC layer was washed in sterile PBS and further centrifuged for 10 minutes at 650 G. The supernatant was discarded and the PBMC pellet was washed again in sterile PBS and centrifuged at 300 G for 10 minutes, followed by a final suspension in 50 ml of complete Roswell Park Memorial Institute (RPMI) media (supplemented with 10% Foetal Bovine Serum (FBS) and 2% Penicillin/Streptomycin (P/S)) before counting.

After counting, CD14^+^ monocytes were isolated from the suspension using the MagniSort^TM^ Human CD14 Positive Selection Kit (Invitrogen) according to the manufacturer’s protocol. Briefly, cells were washed in MACS buffer (PBS with 2% FBS and 0.4% EDTA) and centrifuged at 300 G for 5 minutes. The resultant pellet was resuspended in MACS buffer at 1 x 10^8^ cells/ml in a FACS tube and incubated with anti-CD14 biotin (10 μl per 100 μl cells) for 10 minutes at room temperature. Cells were then washed in 4 ml MACS buffer and centrifuged at 300 G for 5 minutes. Cells were then re-suspended in MACS buffer and incubated with MagniSort^TM^ positive selection beads (15 μl per 100 μl cells) for 10 minutes at room temperature. The volume in the FACS tube was adjusted to 2.5 ml and the tube was placed inside the magnetic field of a EasySep^TM^ cell sorter magnet (STEMCELL Technologies) for 5 minutes. The negative fraction was discarded by inverting the tube while held inside EasySep^TM^ magnet, and the remaining cells were resuspended in 2.5 ml MACS buffer. The wash step was repeated twice and the positive fraction containing the CD14^+^ monocytes was re-suspended in complete RPMI media and counted.

The magnetically sorted CD14^+^ monocytes were plated at a concentration of 500,000 cells per well of a 24 well plate (1 million cells/ml concentration) in complete RPMI containing macrophage colony stimulating factor (M-CSF) at a final concentration of 50 ng/ml. Cells were kept in culture in a 37°C incubator, with 5% CO_2_ and 19% O_2_ concentrations. CD14^+^/CD11b^+^ blood derived macrophages (BDMs) were obtained after 7 days of culture, with media change every 3 days.

### Flow cytometry

On reaching the end of the experiment, cells were suspended in PBS and collected into FACS tubes at a concentration of 500,000 cells per tube.

### For cell-surface markers

Cells were centrifuged at 300 G for 5 minutes followed by incubation in Fc-γ blocker (in FACS buffer: PBS with 2% FBS) for 10 minutes at 4°C to reduce non-specific binding and background fluorescence. Next, cells were centrifuged at 300 G for 5 minutes followed by incubation in the antibodies of interest (detailed in table 1) made up in FACS buffer at a concentration of 10 µl/ml. Each tube was treated with 100 µl of the antibody cocktail for 30 minutes, in the dark at room temperature. The cells were then washed using FACS buffer and centrifuged at 300 G for 5 minutes and used directly for flow cytometry. If being used the next day, the cells were fixed using 1% paraformaldehyde (PFA).

**Table 1.**
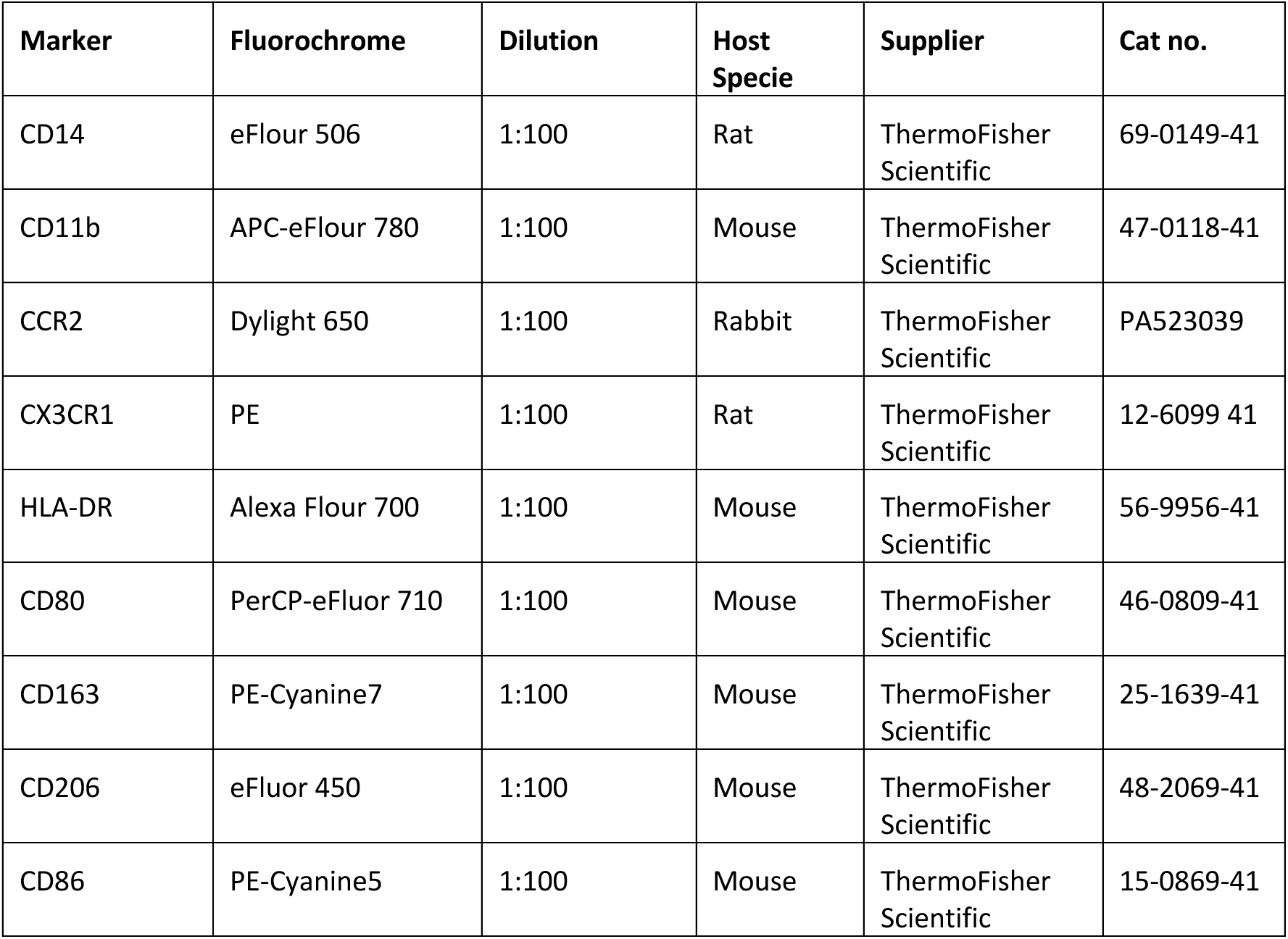
Macrophage-specific antibody list. Macrophage-specific antibodies with fluorochromes and dilution information for flow cytometry.

| Marker | Fluorochrome | Dilution | Host Specie | Supplier | Cat no. |
| --- | --- | --- | --- | --- | --- |
| CD14 | eFlour 506 | 1:100 | Rat | ThermoFisher Scientific | 69-0149-41 |
| CD11b | APC-eFlour 780 | 1:100 | Mouse | ThermoFisher Scientific | 47-0118-41 |
| CCR2 | Dylight 650 | 1:100 | Rabbit | ThermoFisher Scientific | PA523039 |
| CX3CR1 | PE | 1:100 | Rat | ThermoFisher Scientific | 12-6099 41 |
| HLA-DR | Alexa Flour 700 | 1:100 | Mouse | ThermoFisher Scientific | 56-9956-41 |
| CD80 | PerCP-eFluor 710 | 1:100 | Mouse | ThermoFisher Scientific | 46-0809-41 |
| CD163 | PE-Cyanine7 | 1:100 | Mouse | ThermoFisher Scientific | 25-1639-41 |
| CD206 | eFluor 450 | 1:100 | Mouse | ThermoFisher Scientific | 48-2069-41 |
| CD86 | PE-Cyanine5 | 1:100 | Mouse | ThermoFisher Scientific | 15-0869-41 |

**Table 2.** Cardiomyocyte-specific antibody list. Cardiomyocyte-specific antibodies with fluorochromes and dilution information for flow cytometry.

| Marker | Fluorochrome | Dilution | Host Specie | Supplier | Cat no. |
| --- | --- | --- | --- | --- | --- |
| Cardiac troponin T | VioBlue | 1:50 | Human | Miltenyi Biotec | 130-120-542 |
| MLC2a | APC | 1:50 | Human | Miltenyi Biotec | 130-118-546 |
| MLC2v | PE | 1:50 | Human | Miltenyi Biotec | 130-119-581 |
| CD90 | PE | 1:50 | Human | Miltenyi Biotec | 130-128-255 |

Flow cytometry was performed using LD Fortessa (BD).

### For intracellular markers

Cells were centrifuged at 300 G for 5 minutes and resuspend in 500 µl of 1% PFA. The resuspended cells were incubated in room temperature for 20 minutes. Cells were then centrifuged at 300 x g for 5 minutes and resuspended in 1 ml of Flow Buffer 1 (PBS with 0.5% BSA) followed by centrifugation at 300 G for 5 minutes to remove PFA. Cells were then resuspended in 100 µl of Flow Buffer 2 (PBS with 0.5% BSA and 1% Triton-X), with the appropriate stains of interest for 30 minutes at room temperature in the dark. Cells were resuspended in 1 ml of Flow Buffer 2 and centrifuged at 300 G for 5 minutes before finally resuspending in 300 µl of Flow Buffer 2 to perform flow cytometric analysis. Compensation beads (Miltenyi) were prepared and stained the same way as cells.

### RT-qPCR

Samples were stored in (−80°C) Trizol™ until ready for RNA isolation. To carry out RNA isolation, samples in Trizol™ were thawed on ice before adding chloroform. The samples were then vortexed and rested for 15 minutes to allow phase separation. The samples were centrifuged at 12,000 G for 15 minutes at 4°C. The upper aqueous phase containing the RNA was removed and placed on to fresh RNAse-free tubes. GlycoBlue™ was added to the samples, followed by isopropanol and incubated overnight in -20°C to improve the purity of the RNA. The following day, samples were thawed and centrifuged at 14,000 G for 10 minutes at 4°C. The pellet obtained was washed with 70% ethanol followed by centrifugation at 12,000 G for 10 minutes (4°C). The wash step with ethanol was repeated twice and the pellets were allowed to dry entirely before adding 20 µl of RNAse-free water.

The yield and purity of the RNA was assessed using a NanoDrop™. After calculating the total RNA yield, the amount of RNA to be reverse transcribed was decided (500-1000 ng) and was reverse transcribed using a kit from Applied Biosystems (high-capacity cDNA reverse transcription kit). Samples were reverse transcribed using a Thermocycler (BioRad^TM^), to obtain cDNA. The cDNA was reconstituted at 10 ng/µl concentration in nuclease-free water. PCR plates were prepared using SYBR Green Mastermix (Universal iTaq Syber Green, Biorad^TM^), primers of interest and 10 ng of cDNA per well. Plates were run in an Applied Biosystems Fast 7500 PCR machine and the Ct values were used to make calculations for fold change of mRNA expression, using 2^-ΔΔCT^ method [94], using ACTB as the housekeeping gene.

The primer sequences used are detailed in Table 3.

**Table 3.**
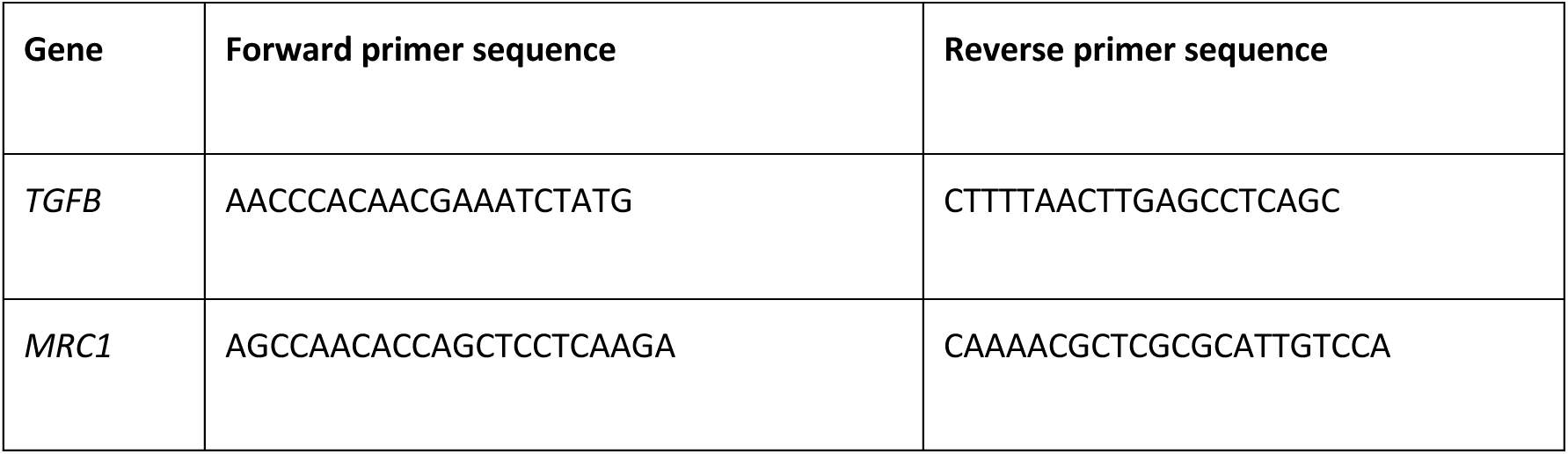
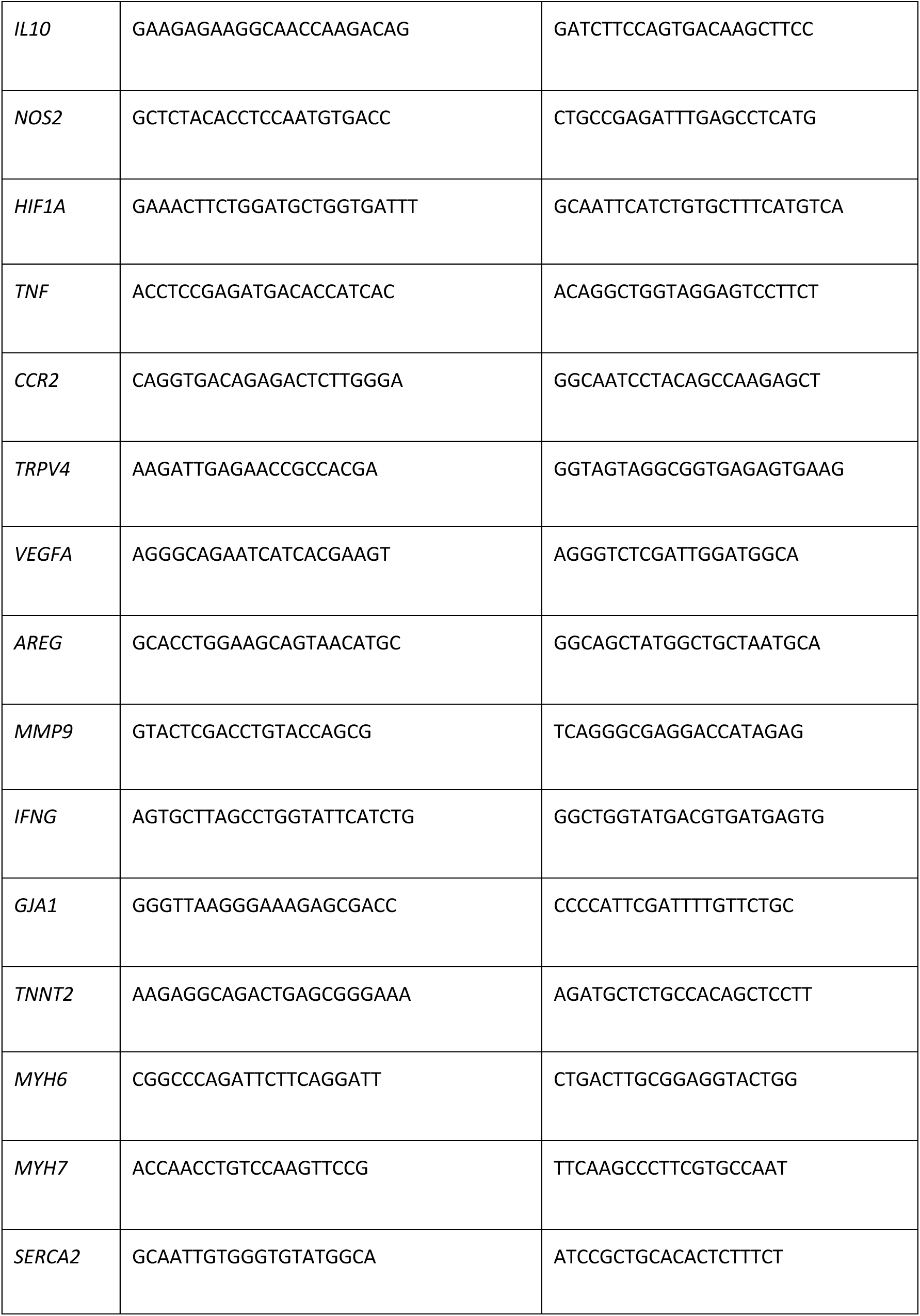
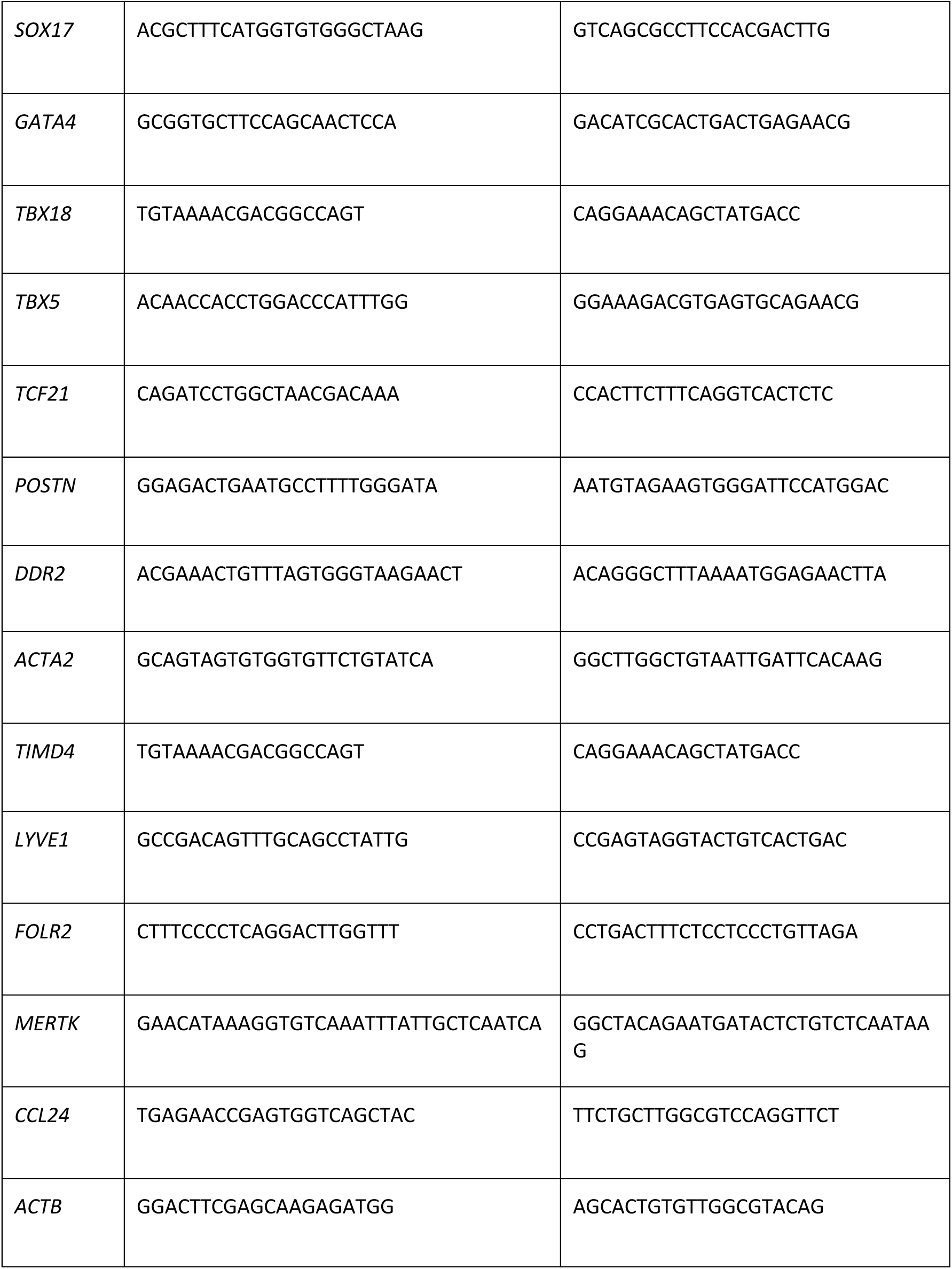
Gene primer sequences.

### ELISA

Supernatants were collected from samples and stored in -80°C for long term storage. ELISA kits (Invitrogen) for TNFα, IL6 and IL10 were used and manufacturer’s instructions were followed. Briefly, high binding plates (Greiner one) were treated with capture antibody diluted in 1x coating buffer and incubated overnight at 4°C. The following day, the unbound capture antibody was removed and the plates were blocked using 1x assay diluent. The plates were incubated in room temperature for 1 hour, after which the samples and standards were added to the plates and incubated overnight at 4°C. The following day, the plates were washed four times followed by an incubation with detection antibody diluted in 1x assay diluent for 2 hours in room temperature. The plates were washed for four times and incubated for 30 minutes with Streptavidin-HRP in the dark. The plates were then washed four times and treated with TMB. The development of the color was stopped by the addition of 1 M sulphuric acid. The absorbance was read at 450 nm using a plate reader.

For measuring exogenous TGF-β1 (Invitrogen), a slightly modified approach was used. After overnight incubation of the plates with the coating buffer, the plates were incubated with the blocking buffer. Samples were prepared by performing a 1:10 dilution in 1x assay buffer, followed by incubation in room temperature for 1 hour with 1 N HCl (20 µl HCl in 200 µl prediluted sample). The samples were then neutralised using 1 N NaOH and added to the wells together with standards and incubated overnight at 4°C. The following day, the plates were washed and incubated with detection antibody diluted in 1x assay diluent for 2 hours in room temperature. The plates were washed for four times and incubated for 30 minutes with Streptavidin-HRP in the dark followed by four more washes before treatment with TMB. The development of the color was stopped by the addition of 1 M sulphuric acid. The absorbance was read at 450 nm using a plate reader.

### Immunofluorescence Imaging

Once the time point was reached, samples were washed with PBS followed by fixation in 4% paraformaldehyde (PFA) (15 minutes for monolayer cells; 30 minutes for hydrogels; 1 hour for human tissues).

#### Sample preparation for 3D samples

Post fixation, samples were incubated in 10% sucrose solution for 2 hours, followed by overnight incubation in 30% sucrose at 4°C. Once the samples settle down at the bottom of the tubes, they were then snap frozen in OCT solution using liquid nitrogen and left at -80°C overnight. Frozen samples were sliced using a cryotome at a thickness of 10 µm and stored at -20°C until ready for staining. Fixed/sliced samples were blocked using a buffer containing PBS with 10% triton-X, 5% Tween-20 and 4% bovine serum albumin (BSA) at room temperature for 30 minutes. The samples were then incubated in the primary antibody of interest, in a buffer containing PBS with 10% triton-X, 5% Tween20 and 1% BSA overnight, at 4°C. The following day, samples were washed 3 times in a wash buffer containing PBS and 0.05% Tween20, followed by incubation in the secondary antibody of choice, for 30 minutes at room temperature. Following one wash with the wash buffer, the samples were incubated with DAPI (1 in 1000 dilution) for 15 minutes at room temperature. Finally, the samples were washed with PBS for 3 times and optionally, incubated overnight in the dark with a drop of prolong gold anti-fade solution. The antibodies used and their respective dilutions as detailed in Table 4.

**Table 4.** Immunofluorescence imaging antibody list. Primary and secondary antibodies with dilution information for immunofluorescence imaging.

| Antibody | Host | Dilution | Supplier; cat no. |
| --- | --- | --- | --- |
| Anti-Cardiac troponin T | Mouse | 1:300 | Abcam; ab8295 |
| Anti-Sarcomeric actinin | Mouse | 1:300 | Fisher Scientific;<br>MA122863 |
| Anti-connexin 43 | Rabbit | 1:300 | Abcam; ab11370 |
| Anti-Rabbit Alexa Flour 594 | Donkey | 1:300 | Abcam; ab150076 |
| Anti-Mouse Alexa Flour 647 | Goat | 1:300 | Abcam; ab314677 |

### Calcium Imaging

iCMs and co-cultures of iCMs and iMacs were cultured in 6 or 24 well plates. On the day of the experiment, 5 mM stock of Cal-520® AM dye was made up to a 3 µM solution, in RPMI media with B-27™ supplement (RPMI+i). Samples were incubated in the media with Cal-520®AM dye for 90 minutes, followed by a media wash and further incubation in RPMI+i. Samples were imaged using a CellSense epi-florescent microscope with heated and motorised stage. Excitation/emission of 490/525 nm was used, and videos were recorded at a fixed exposure time of 100 ms. Videos were analysed using FIJI (ImageJ), where 5 to 9 regions of interest were picked and the intensity variations were tracked. The plots of the intensity variation were used to determine the intensity of calcium signalling (F/F0), beating rate (in beats per minute; bpm), time to peak and time to decay.

### Statistical analysis

Results are represented as mean ± standard deviation from 3 biological replicates, with each point signifying the average of 3 technical replicates. Normalisation tests were performed prior to statistical analysis and comparisons were made based on the experiments and groups being analysed, using GraphPad Prism 10. Kindly refer to figure legends for the exact statistical test performed. * P<0.05, ** P<0.01, *** P<0.001 and **** P<0.0001 were considered significant differences.

## Acknowledgments

The authors acknowledge Dr. Barry Moran, Aoife O’Rourke and Orlaith Walsh from Trinity College Dublin flow cytometry facility and Dr. Gavin McManus from Trinity College Dublin microscopy facility.

## Author contributions

Conceptualisation: MGM, MS

Methodology: MS, LF, JFM, CR

Investigation: MS, SC

Visualisation: MS

Funding acquisition: MGM, MS

Project administration: MGM

Supervision: MGM

Writing – original draft: MGM, MS

Writing – review & editing: MGM, MS, JFM, SC, MB, GF, IT, KDC, LF

## Data and materials availability

All data are available in the main text or the supplementary materials.

## Supplementary Figures

**Supplementary Fig. 1.**
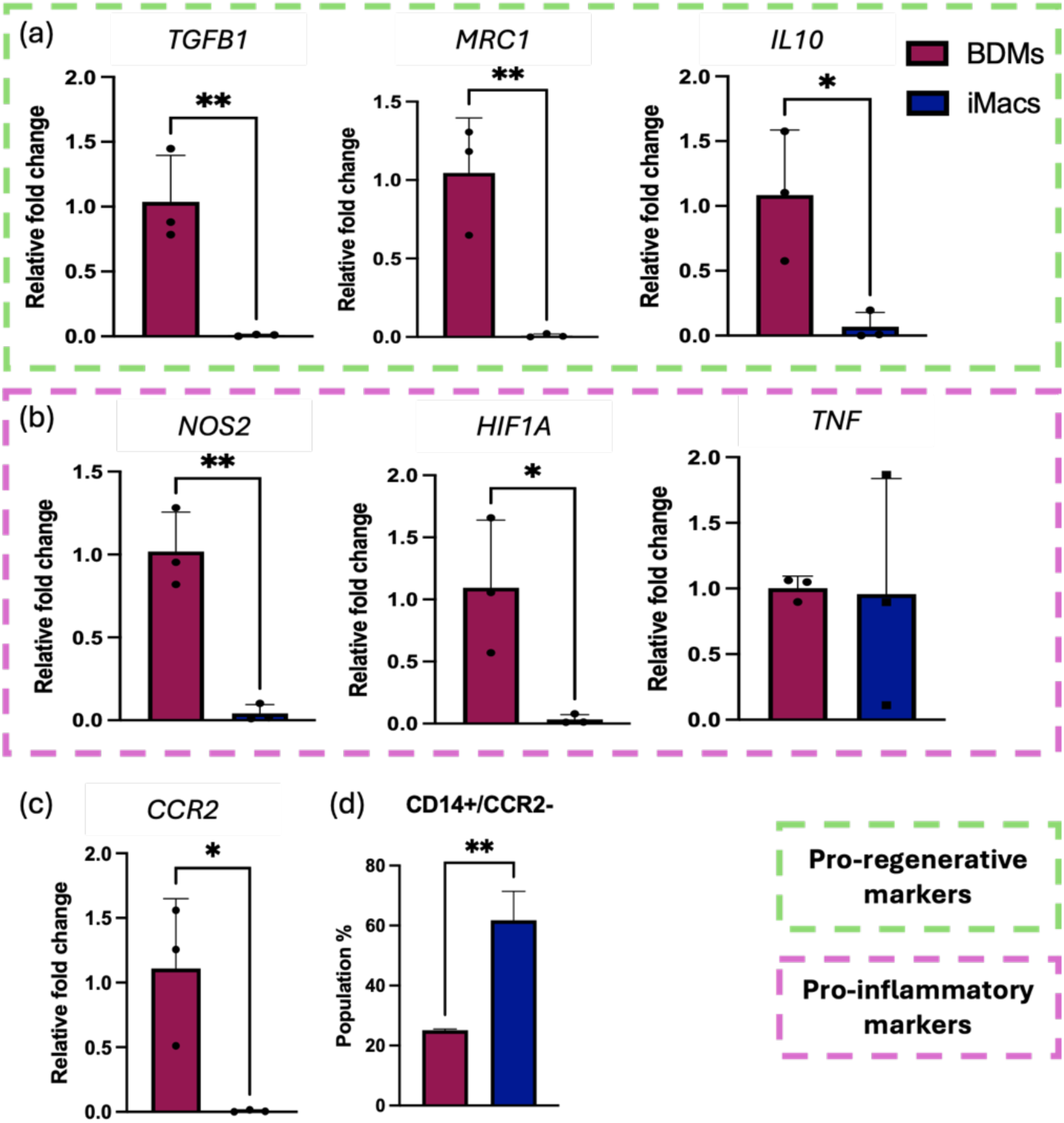
Comparison of the expression of a set of genes between iMacs and BDMs. (**a**) Expression of a set of pro-regenerative genes, (**b**) Expression of a set of pro-inflammatory genes, (**c**) Expression of CCR2 gene, (**d**) Percentage population of cells positive for CD14 and negative for CCR2, using flow cytometry. Statistical analysis - one-way ANOVA with Tukey’s comparison test. * represents P<0.05 and ** represents P<0.01.

**Supplementary Fig. 2.**
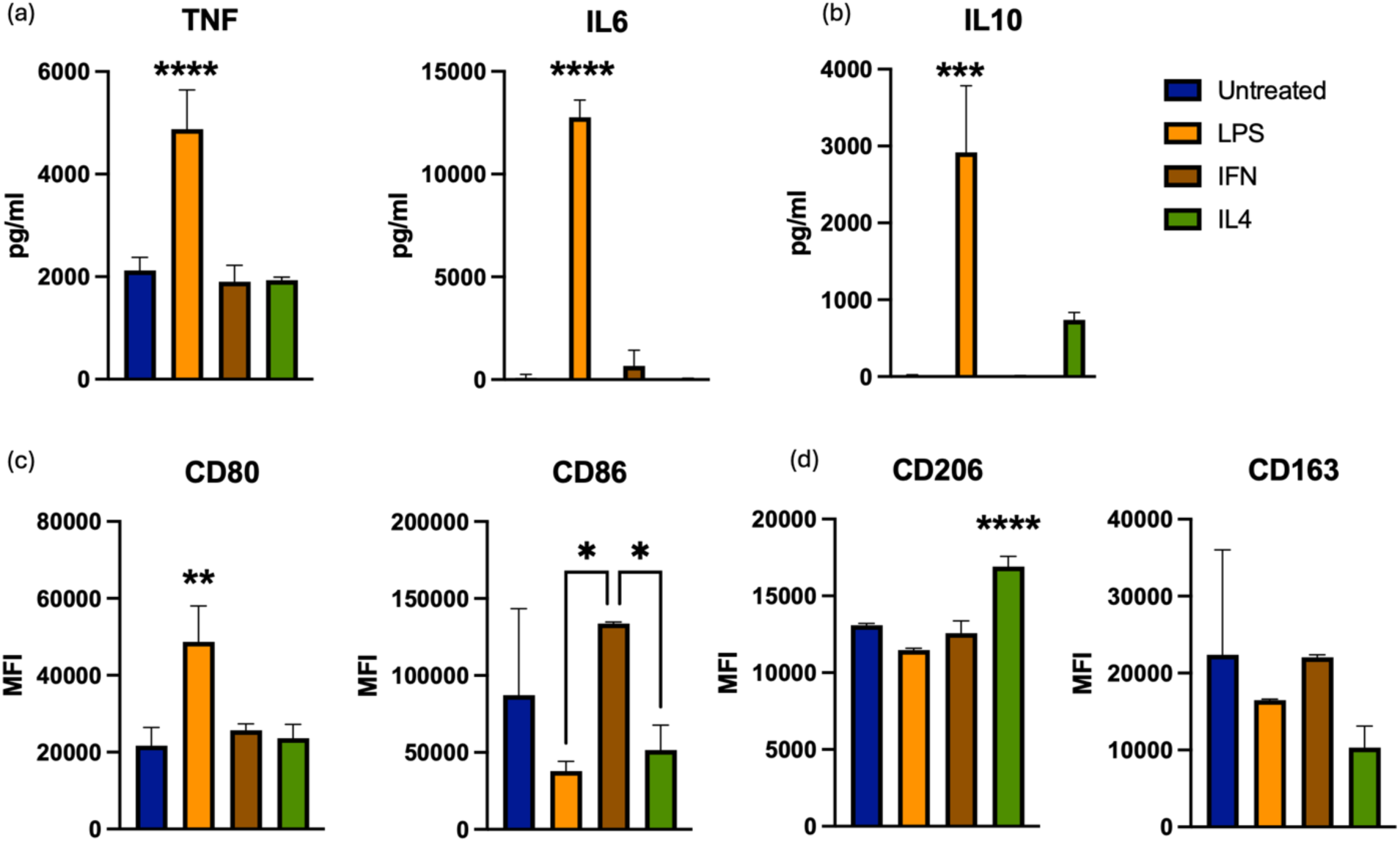
Evaluation of functional ability of iMacs. (**a**) Quantification of release of pro-inflammatory cytokines TNFα and IL6, using ELISA, (**b**) Quantification of release of anti-inflammatory cytokine IL10, using ELISA, (**c**) Quantification of mean fluorescence intensity of pro-inflammatory markers CD80 and CD86, using flow cytometry, (**d**) Quantification of mean fluorescence intensity of anti-inflammatory markers CD206 and CD163, using flow cytometry. Statistical analysis - Ordinary One-way ANOVA. * represents P<0.05, ** represents P<0.01, *** represents P<0.001 and **** represents P<0.0001.

**Supplementary Fig. 3.**
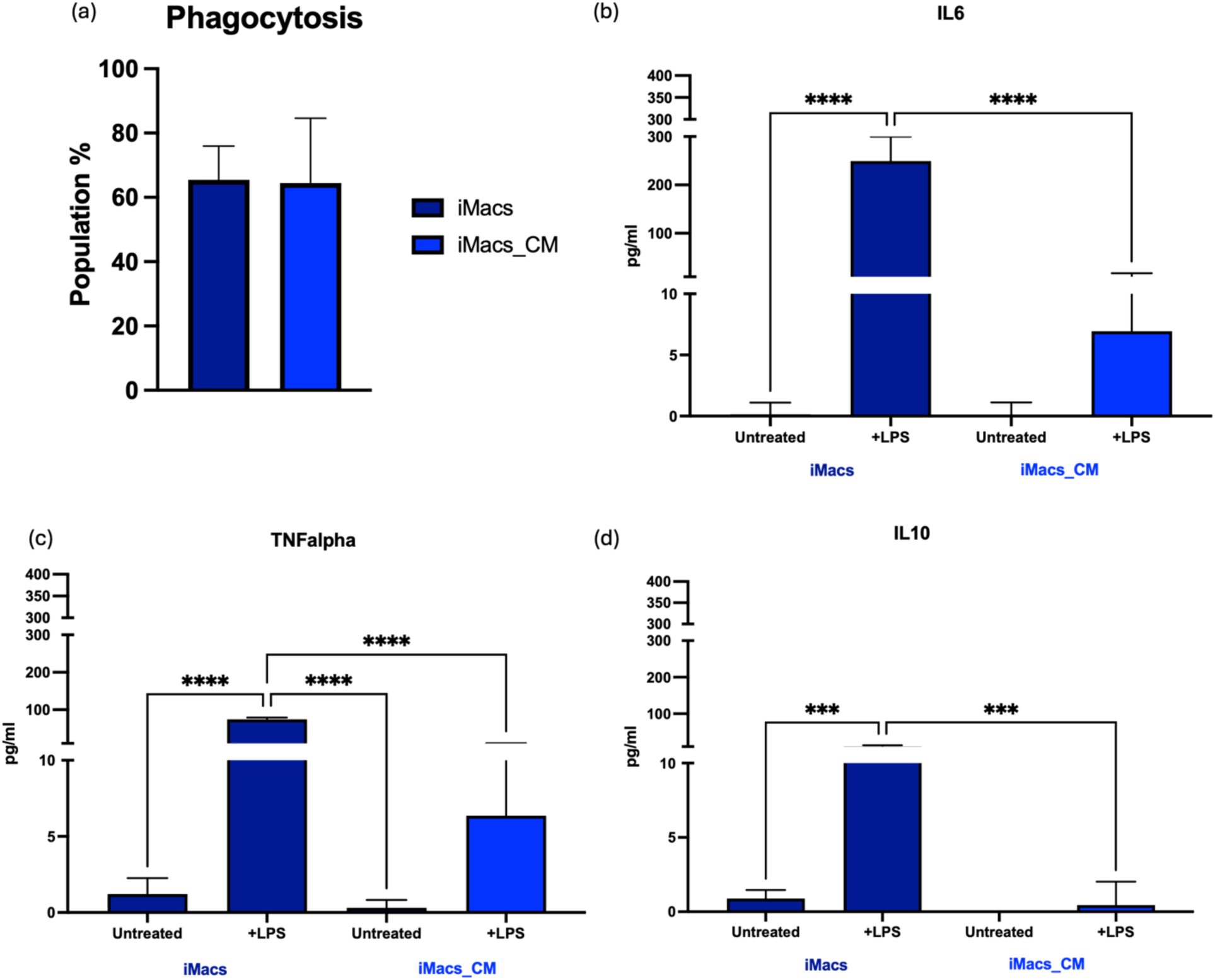
Evaluation of functional ability of iMacs_CM. (**a**) Quantification of phagocytic cell populations in iMacs and iMacs_CM, (**b**) Quantification of IL6 release by iMacs and iMacs_CM, post-LPS stimulation, using ELISA (**c**) Quantification of TNFα release by iMacs and iMacs_CM, post-LPS stimulation, using ELISA (**d**) Quantification of IL10 release by iMacs and iMacs_CM, post-LPS stimulation, using ELISA. Flow cytometry analysis performed using FlowJo. Statistical analysis - Student T test with Welch’s correction for a; Two-way ANOVA with Tukey’s comparison test for b, c, and d. *** represents P<0.001 and **** represents P<0.0001.

**Supplementary Fig. 4.**
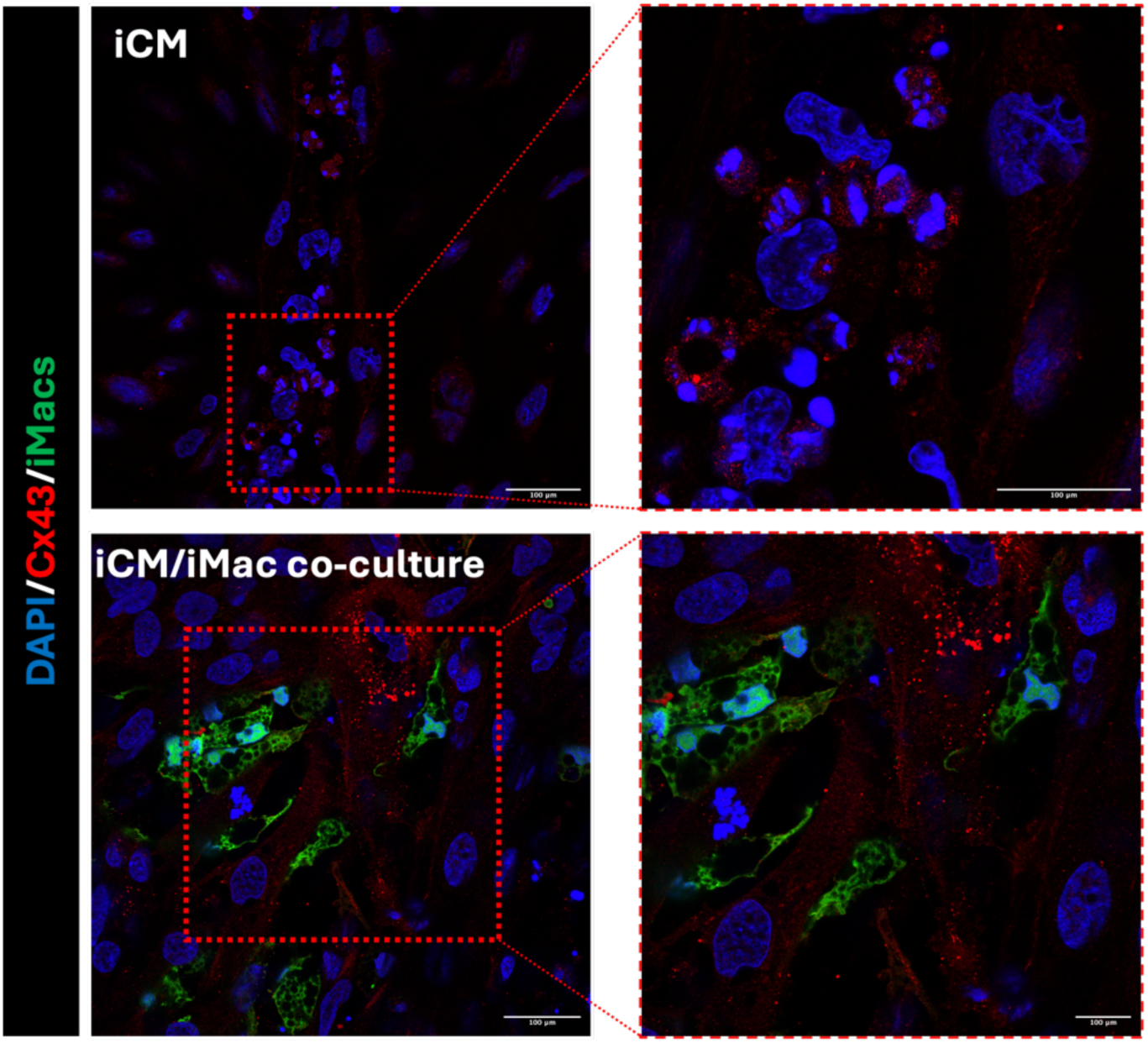
iCMs and iCM/iMacs co-cultures abundantly expressed Cx43. Representative images of iCMs and co-cultures of iMacs and iCMs stained with DAPI and connexin43. iMacs differentiated from SFCi55-ZsGr iPSC line. Scale bar – 100 µm.

**Supplementary Fig. 5.**
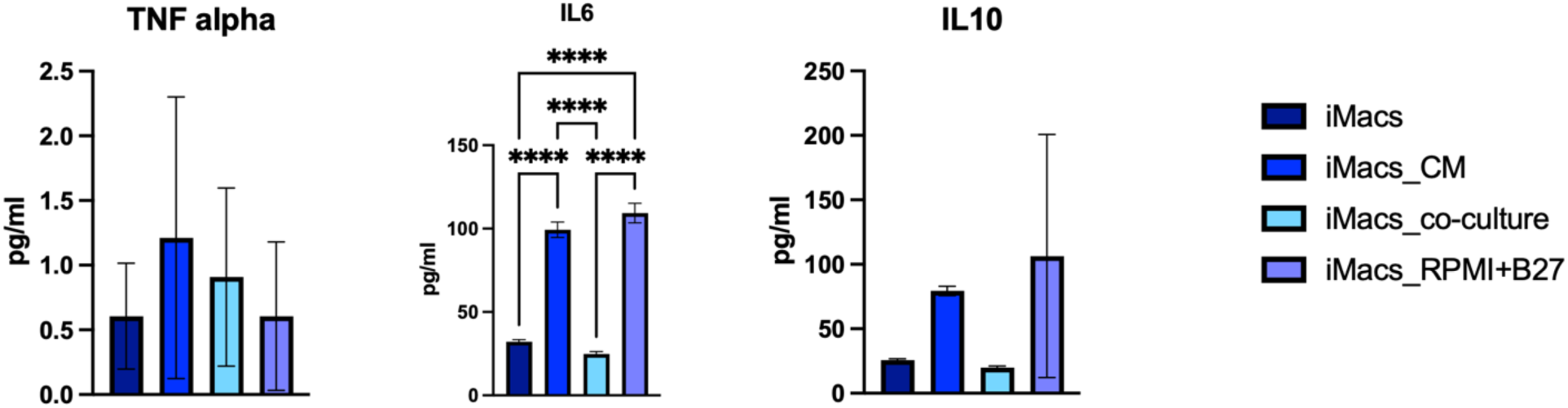
iMacs remain anti-inflammatory when in co-culture with iCMs. Quantification of TNFα, IL6 and IL10 from secretome of iMacs, iMacs_CM, iMacs_co-culture and iMacs cultured in RPMI with B27™, measured using ELISA. Statistical analysis – One way ANOVA with Tukey’s comparison test. **** represents P<0.0001.

**Supplementary Fig. 6.**
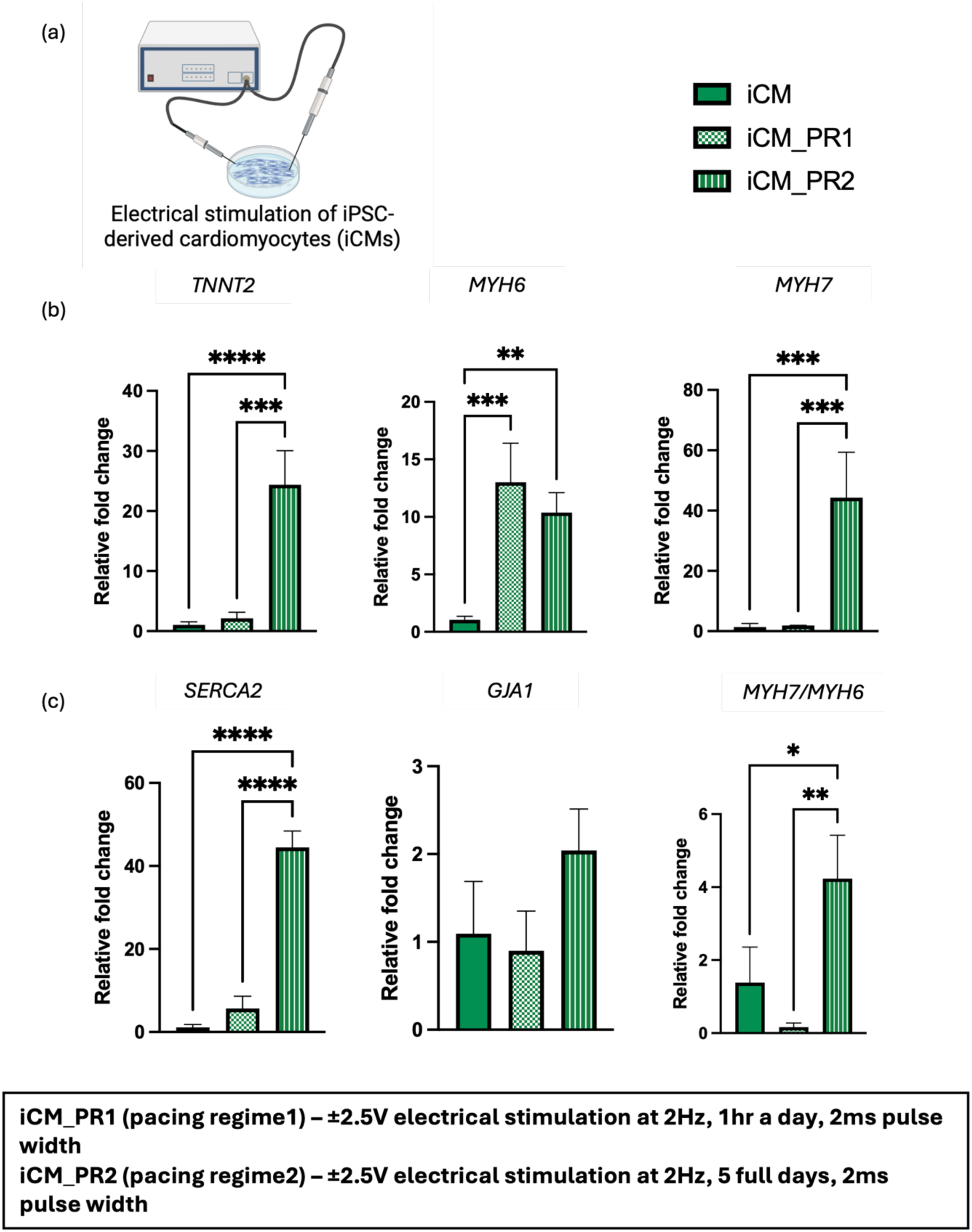
Five days of continuous electrical stimulation improves the expression of maturation genes in iCMs. (**a**) Schematic representation of the experimental set up. (**b**) Expression of the genes TNNT2, MYH6 and MYH7, normalised to unstimulated iCMs. Using qPCR (**c**) Expression of the genes SERCA2, GJA1 and ratio between MYH7/MHY6, normalised to unstimulated iCMs, using qPCR. Statistical analysis – One-way ANOVA with Tukey’s multiple comparison test. * represents P<0.05, ** represents P<0.01, *** represents P<0.001 and **** represents P<0.0001.

**Supplementary Fig. 7.**
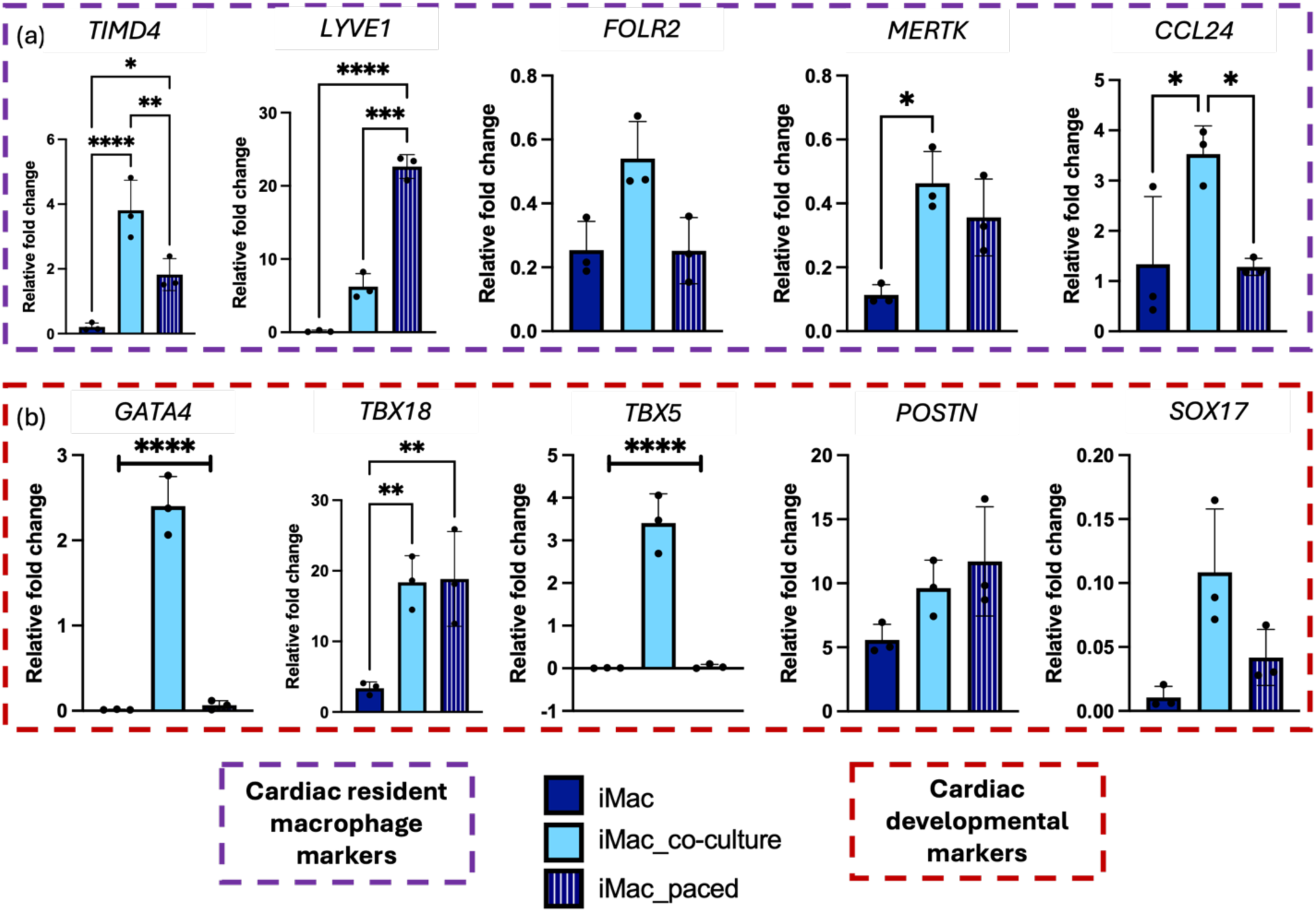
Electrically stimulated iMacs express similar or lower levels of CRM- and cardiac developmental markers, in comparison to iMacs_co-culture. (a) Expression of a set of CRM-specific genes, measured using qPCR. (b) Expression of a set of cardiac developmental genes, measured using qPCR. Statistical analysis - One-way ANOVA with Tukey’s multiple comparison test. * represents P<0.05, ** represents P<0.01, *** represents P<0.001 and **** represents P<0.0001.

**Supplementary Fig. 8.**
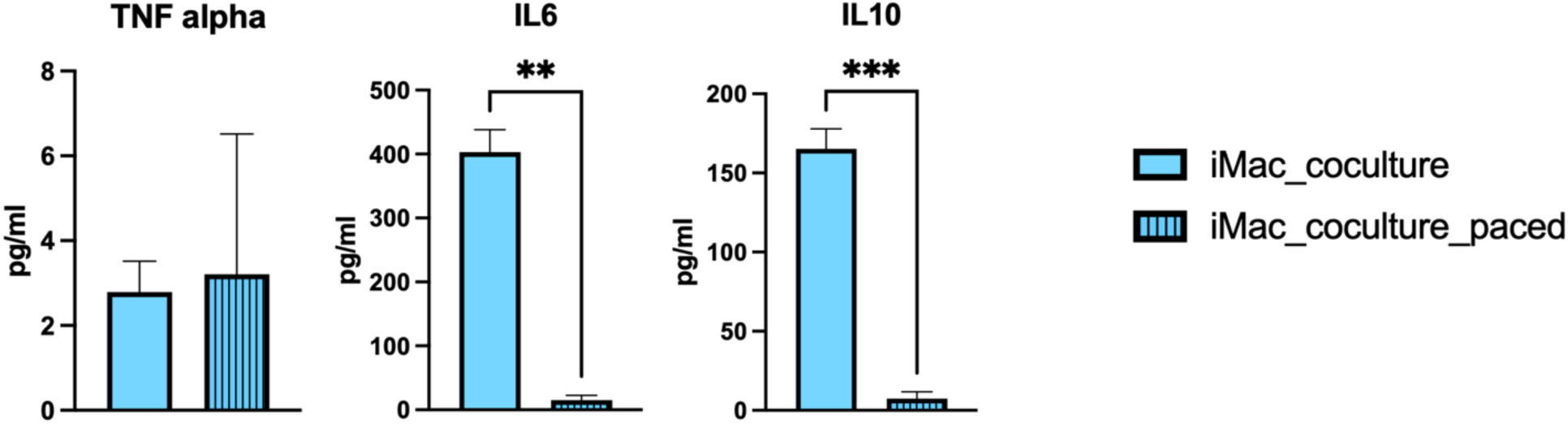
Electrical stimulation does not trigger a pro-inflammatory response in iMacs that are in co-culture with iCMs. Quantification of TNFα, IL6 and IL10 from secretome of iMacs in co-culture with iCMs, with and without ES. Statistical analysis – Student T test with Welch’s correction. ** represents P<0.01 and *** represents P<0.001.

**Supplementary Fig. 9.**
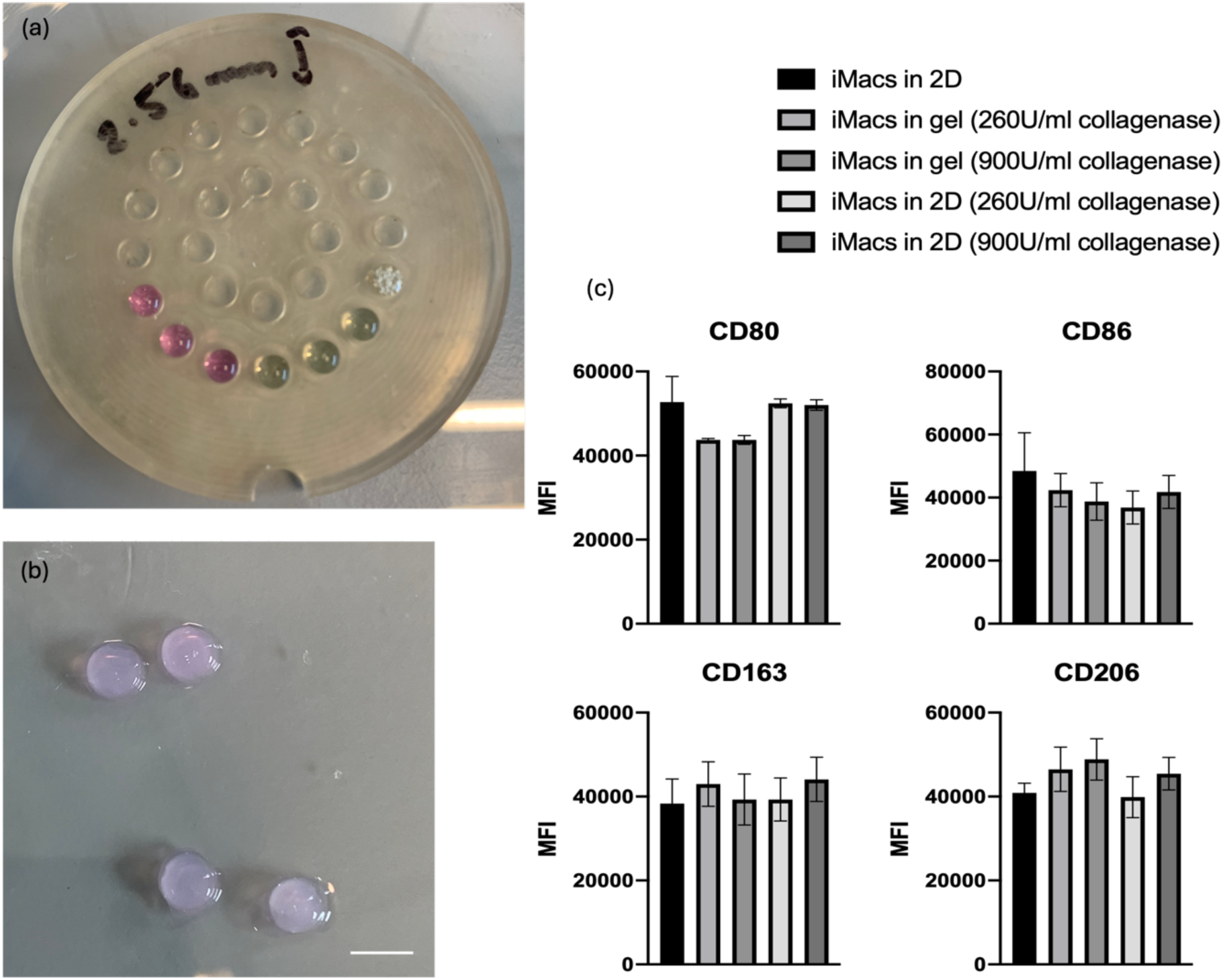
Collagen/Matrigel hydrogel does not trigger an inflammatory response in iMacs. (**a**) Collagen-Matrigel hydrogels with iMacs were casted on PDMS mold of 2.56 mm x 3 mm dimension. (**b**) Representative images of hydrogels after 2 hours of incubation at 37°C. Scale bar – 3 mm. (c) Quantification of the mean fluorescence intensity (MFI) of CD80, CD86, CD163 and CD206 stains on iMacs cultured in Collagen/Matrigel hydrogels for 7 days. Flow cytometry analysis performed using FlowJo. Statistical analysis - One Way ANOVA with Tukey’s multiple comparison test.

**Supplementary Fig. 10.**
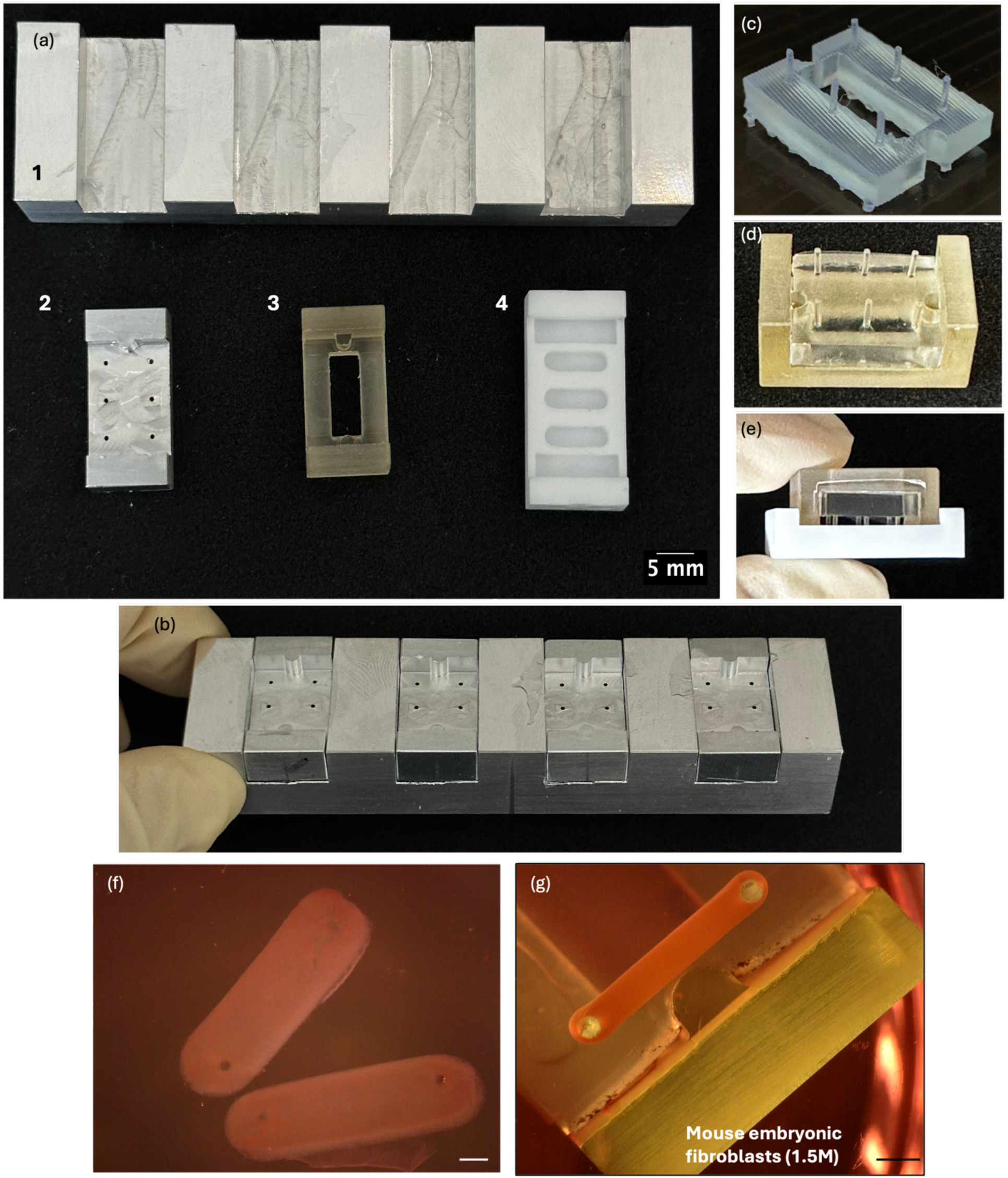
Engineered cardiac tissue fabrication. (**a**) The four components required for the fabrication of ECTs. 1 and 2 are made with aluminium and are attached with each other; 3 is the frame and 4 is the baseplate. (**b**) Parts 1 and 2 attached to each other. (**c**) ECT bioreactor with posts, obtained by casting PDMS on to the aluminium moulds in (b). (**d**) ECT bioreactor attached to the frame. (**e**) Final ECT bioreactor set up with the baseplate. (**f**) Sample image of hydrogels made with the ECT bioreactor (without cells). Scale bar – 1mm. (**G**) Sample image of a tissue made with 1.5 million mouse embryonic fibroblasts, attached to the ECT bioreactor posts. Scale bar – 1mm.

**Supplementary video 1. iMacs integrate well into iCM cultures.** Representative video of iCMs and iMacs (in green) in co-culture, highlighting iMacs preferentially attaching to beating patches of iCMs. iMacs differentiated from SFCi55-ZsGr iPSC line.

**Supplementary video 2. Calcium transients in iCMs.** Representative video of calcium transients in iCMs stained with Cal-520® AM dye, indicating poor synchrony in beating across multiple beating patches.

**Supplementary video 3. Calcium transients in iCM/iMacs co-cultures.** Representative video of calcium transients in iCM/iMacs co-cultures stained with Cal-520® AM dye, highlighting synchronous beating across multiple beating patches of iCMs, facilitated through electrical coupling with iMacs.

**Supplementary video 4. Calcium transients in iCMs cultured with M-CSF.** Representative video of calcium transients in iCMs cultured with M-CSF stained with Cal-520® AM dye, indicating poor synchrony in beating across multiple beating patches.

**Supplementary video 5. Calcium transients in iCMs cultured with iMacs conditioned media.** Representative video of calcium transients in iCMs cultured with iMacs conditioned media stained with Cal-520® AM dye, indicating poor synchrony in beating across multiple beating patches.

